# Protease-mediated degradation of the master transcription factor controls quorum sensing-state transitions in *Vibrio*

**DOI:** 10.1101/2024.02.15.580527

**Authors:** Tanmaya A. Rasal, Caleb P. Mallery, Biqing Liang, Matthew W. Brockley, Chelsea A. Simpson, Abigail D. Padgett, Finley J. Andrew, Laura C. Brown, Jon E. Paczkowski, Julia C. van Kessel

## Abstract

In *Vibrio* species, quorum sensing signaling culminates in the production of the master transcription factor SmcR that regulates group behavior genes in a density-dependent manner. Previously, we identified a small molecule thiophenesulfonamide inhibitor called PTSP that targets the SmcR family of proteins and blocks activity *in vivo*. Here, we used structure-function analyses to identify eight PTSP-interacting residues in the ligand binding pocket that are required for PTSP inhibition of *Vibrio vulnificus* SmcR. Binding of PTSP to SmcR drives allosteric unfolding of the N-terminal DNA-binding domain and, in this state, SmcR is degraded by the ClpAP protease. SmcR degradation controls the timing of the phenotypic switch between high and low cell density, and strains expressing degradation-resistant *smcR* alleles are impervious to changes in cell density state. These studies implicate ligand binding as a mediator of SmcR protein stability and function, which dictates the timing of quorum sensing gene expression in three *Vibrio* pathogens.

**Significance Statement:** SmcR family proteins were discovered in the 1990s as central regulators of quorum sensing gene expression and later discovered to be conserved in all studied *Vibrio* species. SmcR homologs regulate a wide range of genes involved in pathogenesis, including but not limited to genes involved in biofilm production and toxin secretion. As archetypal members of the broad class of TetR-type transcription factors, each SmcR type protein has a predicted ligand binding pocket. However, no ligand has been identified for these proteins that control their function as regulators. Here, we used SmcR-specific chemical inhibitors to determine that ligand binding drives proteolytic degradation *in vivo*, the first demonstration of SmcR function connected to ligand binding for this historical protein family.

## Introduction

Among the >100 species of *Vibrionaceae* that inhabit marine environments, *Vibrio vulnificus* is one of the most potent human pathogens with a mortality rate of >50% in primary septicemia infections (1–3). Historically, infections were endemic to equatorial regions but due to a global rise in temperatures, there has been an alarming increase and spread of vibriosis to regions beyond their typical equatorial habitats (2–4). This has consequently caused worldwide harm to aquaculture industries, natural marine ecosystems, and human health (7–10). Additionally, *Vibrio* species have been shown to evolve by gaining fitness traits through horizontal gene transfer (HGT), including genes encoding multidrug resistance (9, 10). The emergence of multidrug resistant *Vibrio* strains has prompted a need to develop alternative strategies to treat vibriosis.

In *Vibrio* species, one major regulatory system that controls expression of virulence genes is quorum sensing (QS), a cell-to-cell signaling system that allows bacteria to control group behaviors in response to alterations in population density (11–13). The model presented in **Figure S1** depicts the QS signaling system defined in *Vibrio campbellii*. At low cell density (LCD), autoinducer concentrations are low, and multiple unbound receptors act as kinases to phosphorylate LuxU, the phosphotransfer protein. LuxU phosphorylates the response regulator LuxO, and phosphorylated LuxO, together with σ-54, activates expression of the five Qrr small RNAs (16). The Qrrs repress the master QS transcription factor, a family of proteins only found in *Vibrios* and collectively called the LuxR/HapR family (17). The Qrrs also activate translation of the LCD regulator AphA, leading to expression of genes for individual behaviors (16–18). At high cell density (HCD), binding of autoinducers and/or nitric oxide (19, 20) to the receptors changes their activities to those of phosphatases, resulting in dephosphorylated LuxO and loss of Qrr expression. Thus, production of LuxR/HapR is highest at HCD, and this master transcription factor regulates the expression of genes encoding for proteins with various functions (19, 20, 23). The master QS transcriptional regulators in each *Vibrio* species, *e.g*., *V. campbellii* LuxR, *V. cholerae* HapR, *V. parahaemolyticus* OpaR, and *V. vulnificus* SmcR, are highly conserved in pathogenic *Vibrios* and are important regulators of virulence gene expression (17, 24). These proteins are not structurally or functionally the same as the LuxR-type proteins found associated with LuxI acyl-homoserine lactone synthases in other Gram-negative bacterial QS systems (12, 23, 24). Rather, the master QS transcription factors that we discuss here called the LuxR/HapR family are part of the broad TetR family of proteins. TetR proteins are characterized by homodimeric complexes with an N-terminal helix-turn-helix DNA binding domain (DBD) and a C-terminal dimerization domain with a putative ligand-binding pocket (LBP), which typically mediates the conformation of the DBD. Thus, TetR DNA-binding activity is classically controlled in a ligand-dependent manner (27). Crystal structures of SmcR and HapR revealed a putative LBP near the dimerization domain that is characterized by hydrophobic residues and is hypothesized to contain an unidentified native ligand (23, 24). However, no ligand has been identified to date. Thus, functional studies of the LuxR/HapR family of proteins and their ligand binding partners have been limited.

Because LuxR/HapR proteins are the central regulators of QS signaling genes, they have been a focus of research since their discovery (28). Indeed, their importance in *Vibrio* virulence gene regulation is well-documented; deletion of *smcR* or inhibition of SmcR in *V. vulnificus* decreases pathogenesis in both mouse and shrimp models (29, 30). Recent studies have shown that QS inhibitors are feasible alternatives to traditional antibiotics with demonstrated efficacy in infection models (31, 32). Because these inhibitors do not affect bacterial viability, they are hypothesized to generate less selective pressure for evolving resistance (33, 34). LuxR/HapR-specific inhibitors have been identified via *in vitro* and *in silico* screening efforts and include molecules with functional groups such as aromatic sulfonamides, enones, sulfones, furanones, brominated thiophenones, and cinnamaldehydes (35–41). Each of these inhibits QS phenotypes to some degree in *V. campbellii* and/or other *Vibrio* species. However, most of these either did not result in large changes in QS or required high concentrations to be effective. Kim *et. al.* identified a selective *Vibrio* QS inhibitor named Qstatin [1-(5-bromothiophene-2-sulfonyl)-1H-pyrazole], which inhibits *V. vulnificus* SmcR (29). Based on biochemical assays, they hypothesized that Qstatin binds to the putative LBP in SmcR, changing the entropic and enthalpic components of the system and altering SmcR regulatory activity *in vivo*. Their data showed that Qstatin resulted in SmcR dysfunction, eliminating the expression of the SmcR regulon required for virulence, biofilm dynamics, and motility/chemotaxis. Qstatin markedly attenuated different representative QS-regulated phenotypes in the test *Vibrio* species, including virulence against the brine shrimp (*Artemia franciscana*) (29). However, the mechanism of action of Qstatin was not clear; addition of Qstatin did not alter DNA binding more than 2-fold *in vitro* (29), and that result was not observed in DNA binding assays performed by our group (42).

Subsequently, our group synthesized a panel of thiophenesulfonamide inhibitors and screened them for their inhibitory activity against different LuxR/HapR proteins. We identified several compounds that displayed equal or better efficacy compared to Qstatin against LuxR/HapR homologs in various *Vibrio* species with no impact on bacterial viability (43). From that panel, PTSP (3-**<u>p</u>**henyl-1-(**t**hiophen-2-yl**<u>s</u>**ulfonyl)-1*H*-**<u>p</u>**yrazole) was the most effective inhibitor (**Fig. 1A**). It is structurally different from Qstatin such that it has an additional phenyl group on the pyrazole ring of the molecule and no bromine on the thiophene ring. It inhibits LuxR/HapR homologs in different *Vibrio* species at nanomolar to micromolar concentrations with distinct efficacies based on their respective putative LBP (41). PTSP shows the strongest inhibition of *V. vulnificus* SmcR but does not inhibit *V. cholerae* HapR. Based on these results, the Raychaudhuri lab sought to determine the molecular basis of HapR insensitivity to Qstatin and its derivatives (44). They constructed a variant HapR by replacing four native LBP residues with SmcR residues, two of which our group had shown to be necessary for SmcR sensitivity to PTSP. They made single, double, triple, and quadruple substitution mutants at residues Y76F, L97I, I141V and F171C (43, 44). Their assays showed that only the quadruple mutant was susceptible to Qstatin and its phenyl derivative inhibitor called IMT-VC-212 (44).

**Figure 1.**
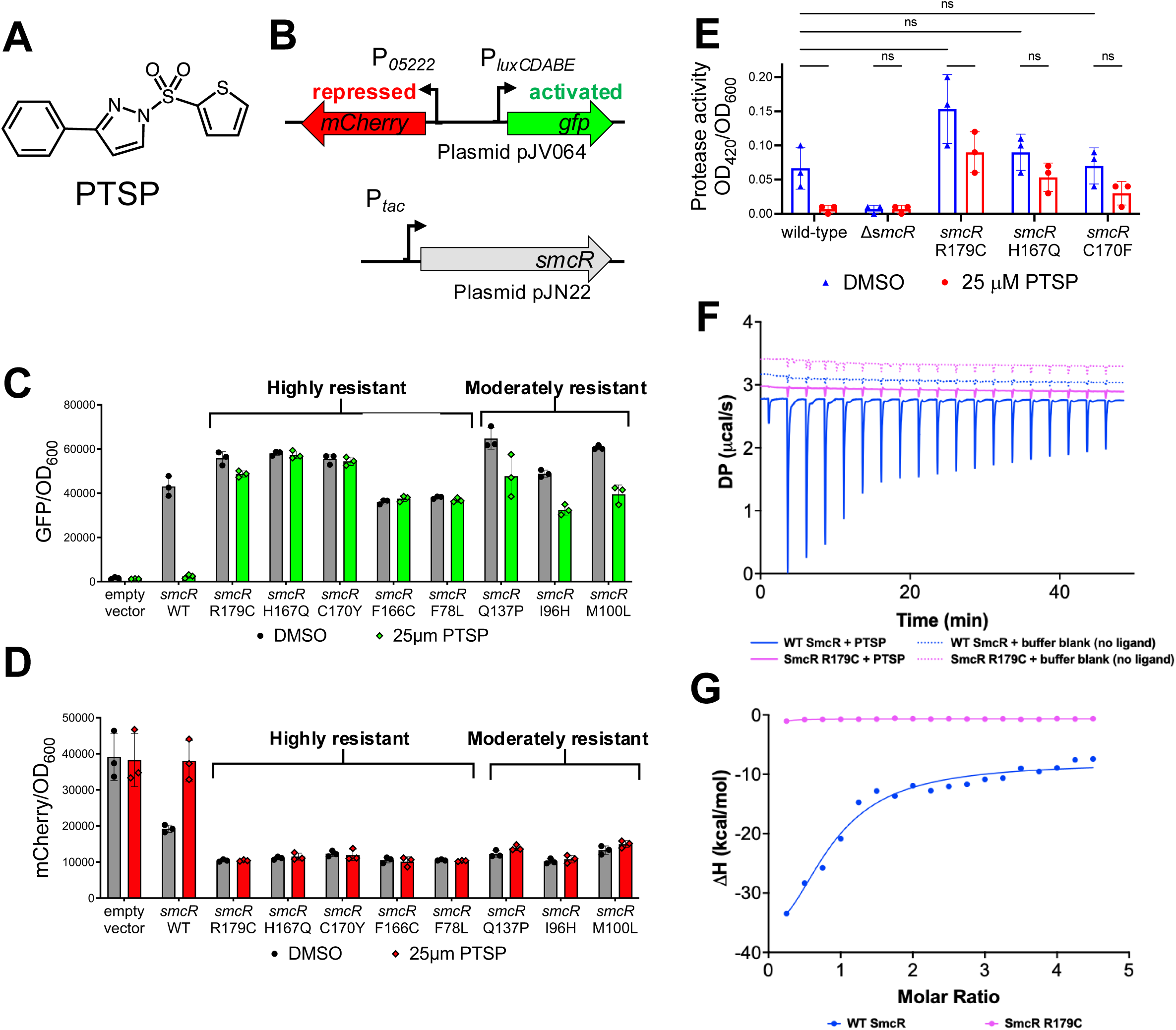
Forward genetic screen identifies mutant *smcR* alleles resistant to PTSP. (A) Structure of PTSP compound. (B) Diagram of the two-plasmid screen in *E. coli* in which wild-type or mutant *smcR* alleles encoded on plasmid pJN22 were screened for activity using a two-color reporter plasmid pJV064. The *luxCDABE* promoter is activated by SmcR and the *05222* promoter is repressed by SmcR. (C, D) *E. coli* strains contained plasmids pJN22 (wild-type *smcR* allele) or mutant pJN22 plasmids from library screening and pJV064. SmcR expression was induced with 50 μM IPTG and cultures treated with either 25 μM PTSP or an equal volume of DMSO solvent. GFP (C) and mCherry (D) relative to the OD_600_ were measured and plotted. (E) Protease activity (OD_420_/OD_600_) of wild-type and mutant *V. vulnificus* strains. (F) Representative isothermal titration calorimetry of WT SmcR (blue) and SmcR R179C (magenta) with (solid lines) or without (dotted lines) PTSP. (G) Transformed data from (F). DP and ΔH denote differential power and enthalpy, respectively. Statistical analyses: a two-way analysis of variance (ANOVA) was performed on normally distributed data (Shapiro-Wilk test; *n*=3; **, *p*=0.01; ***, *p*=0.001).

Here, we used a genetic screen and biophysical techniques to determine PTSP interactions with SmcR. Our results show that multiple residues in the LBP of SmcR are critical for PTSP binding *in vivo*. Substitutions at residues in the LBP abrogate PTSP inhibition and restore wild-type activity to SmcR *in vivo*. PTSP binding to SmcR increases protein degradation by the ClpAP protease *in vivo.* Clp proteases belong to the highly conserved ATP-dependent proteases- these are composed of a proteolytic core (ClpP) and one or more unfoldases (ClpA, ClpC, ClpX, etc.). They play a crucial role in protein quality control and removal of damaged or misfolded proteins (45, 46). We also observed that SmcR variants that are resistant to PTSP were not susceptible to this degradation. Thus, the mechanism of thiophenesulfonamide inhibition of SmcR is driven by proteolysis *in vivo*, indicating that the stability of SmcR, and possibly its conformation, is connected to interactions with the compound in its LBP. This mechanism was observed in multiple *Vibrio* species. Moreover, ClpAP-mediated turnover of SmcR impacts the transitions between LCD and HCD signaling states, broadly implicating the importance of SmcR ligand binding in *Vibrio* signaling and response.

## Results

### A forward genetic screen identifies *smcR* alleles resistant to PTSP

Our previous study showed that substitutions F75Y or C170F rendered SmcR resistant to PTSP, whereas I96L and V140I did not affect PTSP sensitivity nor function (43). However, this was a limited reverse genetics experiment that encompassed only four substitutions in the putative LBP and did not determine if other residues are also involved in binding to ligand. To determine which SmcR residues are important for PTSP binding, we used a forward genetics approach in which we generated a *smcR* mutant library using random mutagenesis by error-prone PCR. The mutant libraries were assayed in a previously established dual-reporter bioassay in *E. coli* that contains *V. campbellii* promoters activated or repressed by LuxR/HapR-type proteins: 1) an activated reporter cassette comprising the *luxCDABE* promoter driving *gfp* expression, and 2) a repressed reporter cassette comprising the *05222* promoter driving *mCherry* expression (**Fig. 1B**) (19). Thus, in cells expressing wild-type *smcR*, GFP expression is high and mCherry expression is low. Conversely, in *smcR*-expressing cells in the presence of PTSP, GFP expression is low and mCherry expression is high, mimicking cells that do not express *smcR* (**Fig. 1C, 1D**). As with our previous reverse genetics assay, we sought to identify mutant *smcR* alleles that were resistant to PTSP and expressed GFP and mCherry at levels comparable to wild-type *smcR* cells in media without PTSP. Using flow cytometry, we sorted PTSP-treated cells based on their GFP and mCherry expression profiles into high and moderate resistance classes based on net difference in GFP values compared to DMSO-treated cells. (**Fig. 1C, 1D**). From screening ∼60,000 mutant alleles, we identified multiple isolates with mutant *smcR* alleles with substitutions at residues F78, I96, M100, Q137, F166, H167, C170, and R179 (**Fig. 1C, 1D, Fig. S2**).

To assess the effect of substitutions in SmcR on QS regulation and growth, we focused on three PTSP-resistant *smcR* alleles with substitutions at H167, C170 and R179 and constructed isogenic mutant *V. vulnificus* strains. *V. vulnificus* utilizes various virulence factors, such as exotoxins and other biofilm-related factors for its persistence in the environment and pathogenesis during infection (43). We examined the effect of PTSP on protease activity. The *vvpE* gene encodes a protease, and this gene is known to be activated at HCD by SmcR activation (48). We hypothesized that the resistant mutant strains would exhibit no change in protease activity in the presence of PTSP but expected protease activity to decrease or be eliminated upon PTSP binding in the wild-type strain. First, we qualitatively measured each strain for growth and observed that the mutations did not lead to significant growth attenuation either in the presence or absence of PTSP (**Fig. S3**). We next measured protease activity in *V. vulnificus* supernatants using an azocasein-based assay. We observed that protease activity in wild-type *V. vulnificus* decreased ∼10-fold upon treatment with 25 µm PTSP (**Fig. 1E**). Mutant strains treated with PTSP exhibited similar protease activities to the untreated cultures; although there were slight decreases in protease activity for each mutant, the only significant difference was in the strain expressing the SmcR variant R179C. We note that the protease activity of this mutant strain is significantly increased compared to wild-type in the untreated cultures, indicating that the R179C substitution somehow alters basal SmcR activity, a point we return to below. From these data, we conclude that we have identified mutants of *smcR* that render SmcR resistant to PTSP and maintain and/or increase SmcR activity *in vivo*.

Previously, we had shown that PTSP binds *V. vulnificus* SmcR *in vitro* with greater affinity compared to *V. campbellii* LuxR (43). To determine if SmcR R179C could bind PTSP, we analyzed the binding affinity of PTSP to purified wild-type SmcR and the SmcR R179C variant by isothermal titration calorimetry (ITC). In its dimeric form, SmcR has two identical putative ligand-binding pockets. We incubated 10 µM of PTSP in the cell and titrated 100 µM of wild-type SmcR or SmcR R179C over 18 consecutive injections (**Fig. 1F, Fig. S4**). We found that the wild-type SmcR-PTSP interaction had a K_d_ of approximately 1 µM, while the SmcR R179C-PTSP interaction was sufficiently weak enough that a K_d_ could not be calculated (**Fig. 1G, Fig. S4**). These findings are consistent with the *in vivo* reporter data that showed that the SmcR R179C variant was recalcitrant to PTSP inhibition (**Fig. 1C**), further validating our structural and genetic approaches. These data suggest that PTSP binds SmcR specifically with ∼1 µM affinity, and this interaction is disrupted in LBP variants of SmcR as observed with R179C.

### PTSP binds to SmcR in the putative LBP

We used X-ray crystallography to determine the structural basis for the SmcR-PTSP interaction. We note that initial attempts to soak PTSP into already-formed SmcR protein crystals did not result in electron density maps that contained density consistent with a bound PTSP molecule, which was the method performed for the SmcR-Qstatin structure published (29). Instead, we added PTSP at the time of protein induction and screened for conditions that could promote crystallization of SmcR pre-bound to PTSP. We determined the structure of full-length SmcR bound to PTSP at 2.1 Å using molecular replacement (**Fig. 2A**; **Table S1; Movie S1**). The N-terminal DBD was largely disordered and residues 1- 15 could not be modeled with high confidence and, thus, were not included in our final model (**Fig. S5A**). Additionally, the B-factors were high for residues in the more ordered region of the N-terminus (residues 16-62) that could be modeled compared to the rest of the structure (**Fig. S5A**). Indeed, it is possible this region of protein is actively unfolded or destabilized by the presence of PTSP, a point we address in our biochemical analyses of the SmcR-PTSP complex below. Conversely, the C-terminal domain, which contains the putative LBP, was well-ordered and allowed for the unambiguous placement of PTSP (**Fig. 2B, Fig. S5B**). The putative LBP is bipartite with distinct hydrophobic and hydrophilic regions, similar to what is observed in the ligand binding domains of LuxR-type proteins (49). One-half of the LBP is comprised of hydrophobic residues (F75, F78, L79, I96, M100, W114, F115, V140, F166, C170), which can accommodate the phenyl and pyrazole rings of PTSP (**Fig. 2B**). Additionally, the region of the LBP adjacent to the solvent accessible channel that allows for ligand accessibility to the binding site is lined with hydrophilic residues, which can act as a shield for the surrounding bulk solvent. Of note, residue R179 from one monomer of SmcR contacts water residues in the accessible ligand channel of the opposing monomer, potentially acting as a gate for ligand binding (**Fig. 2B; Fig. S5C)**. Furthermore, hydrophilic residues (N133, Q137, N164, H167) also contact water molecules and the thiophene head group of PTSP (**Fig. 2B**). In particular, residue N133 forms an important hydrogen bond with the sulfonyl group in PTSP (**Fig. 2B**). Nine of these residues were discovered as being important for binding to PTSP in our forward genetic screen of SmcR or through reverse genetics (43) (**Fig. 1B, 1C, S5**), lending validity to our model. Previous work in our lab revealed that HapR from *V. cholerae* was resistant to PTSP (41). To understand the molecular basis for the selectivity of SmcR for PTSP, we compared the LBP of SmcR to HapR. The most significant difference between the structures is at residues C170 in SmcR and F171 in HapR (**Fig. S5D**). When the PTSP-bound structure of SmcR and the apo structure of HapR were overlayed, residue F171 from HapR overlapped with the phenyl ring in PTSP, revealing a significant steric clash that would prevent PTSP binding to HapR (**Fig. S5D**).

**Figure 2.**
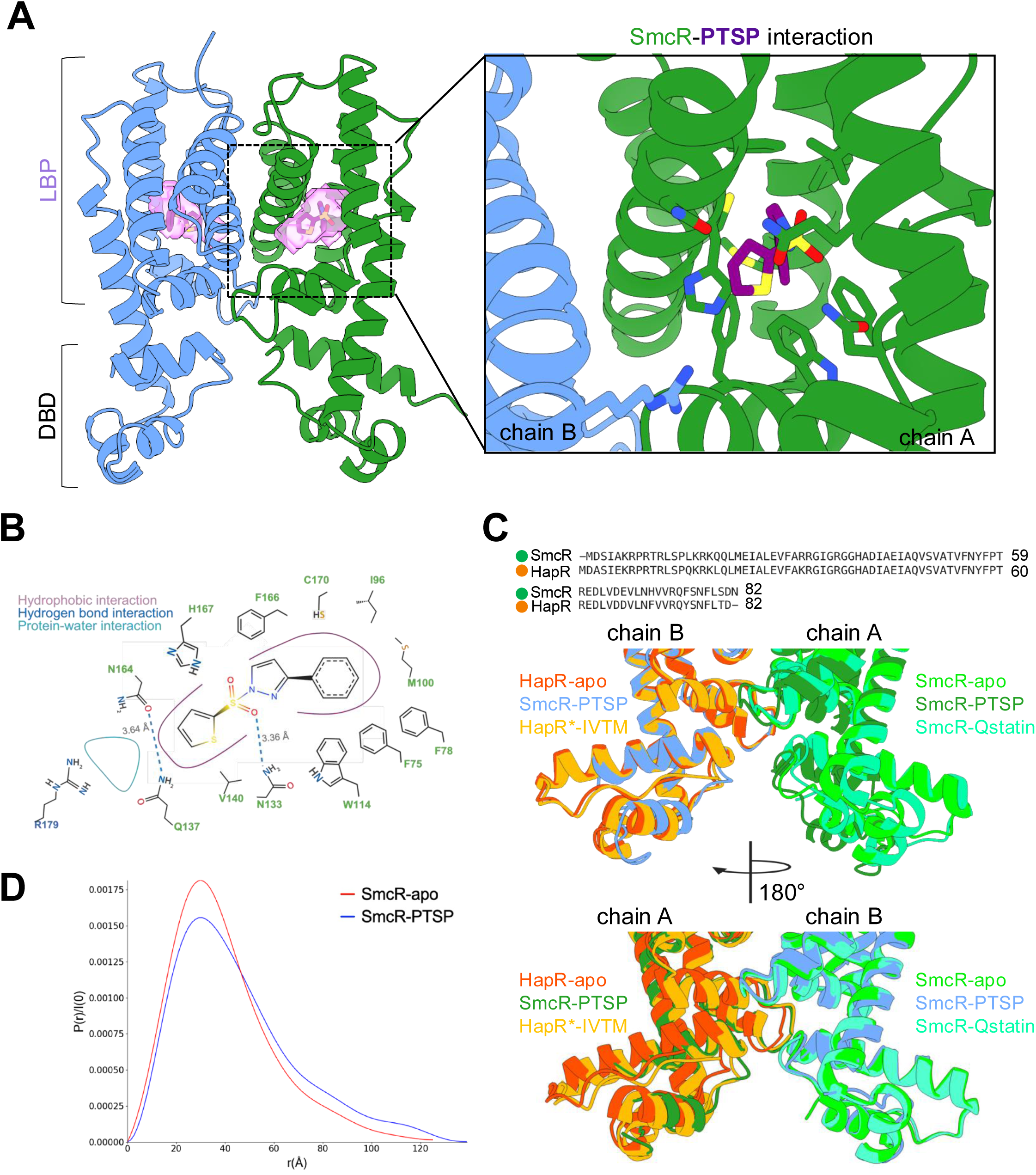
Structural basis for SmcR ligand binding. (A) X-ray crystal structure of homodimeric SmcR (blue/green) bound to PTSP (purple). Solvent accessible ligand binding pocket is shown as a surface in light purple (left). Residues important for contacting PTSP in the SmcR LBP (right). (B) 2D projection of the SmcR-PTSP binding interface showing hydrophobic contacts (purple lines), hydrogen bond interactions (blue), and solvent interactions (cyan). (C) Sequence and structural alignment of HapR and SmcR. The X-ray crystals structures of SmcR-apo (light green; PDB: 6WAE), SmcR-PTSP (colored by chain as in (A)) , SmcR-Qstatin (blue-green; PDB: 5X3R), HapR-apo (orange; 7XXO), and HapR*-ITVM (light orange; PDB: 7XYI). HapR* indicates the HapR quadruple variant from Sen et al. 2023 that stably binds to thiophenesulfonamide inhibitors. (D) The estimated pair-distance distribution function, P(r) normalized by the scattering intensity, I(0). The P(r) function is based on the Indirect Fourier Transform of SAXS data for SmcR-apo (red) and SmcR-PTSP (blue).

In recent years, SmcR and HapR have been targets for small molecule modulation for QS-dependent repression of virulence (26–32, 41). SmcR and HapR variants bound to molecules possessing similar moieties to PTSP have been solved, allowing us to determine the mechanism driving receptor inactivation. In Fig. 2C, we overlayed the structures of SmcR-apo, SmcR-Qstatin, HapR (Y76F, L97I, I141V and F171C)-IMT-VC-212 with a focus on the DBD of the receptors, specifically residue N55 in SmcR (N56 in HapR), which we previously showed to be important for recognizing DNA to initiate transcription (42). Typical of TetR receptors, SmcR and HapR both have an extended α-helix (residues 60-84) that connects the putative LBP and the DBD. In the SmcR-PTSP structure, the first α-helix of the DBD is largely disordered, resulting in a displacement of N55, allowing the residue to point inward toward the core of the DBD rather than the bulk solvent (**Fig. 2C**; **Fig. S5E**). Collectively, from these data, we conclude that PTSP forms specific interactions with the LBP of SmcR, and alterations of the DBD conformation, at the α-carbon backbone level, are observed in the PTSP-bound structure compared to the apo-SmcR structure. However, due to the nature of the disordered N-terminus, we cannot reliably determine the specific molecular interactions that might drive such a change.

### PTSP promotes SmcR flexibility in solution

To assess the potential conformational changes induced by PTSP binding in solution, we performed small-angle X-ray scattering (SAXS) experiments using SmcR-apo and SmcR-PTSP (**Fig. S6 and S7**). Consistent with previous work (42), SAXS data collected for SmcR-apo fit well with high-resolution X-ray structure data (PDB: 6WAE) (low χ^2^, with a value of 0.80. We observed a poor fit of the SmcR-PTSP SAXS data to high-resolution data, with a χ^2^ value of 2.58 (**Fig. S8**, **Table S2**). This observation agrees with our X-ray crystallographic data, in which we observed disordered regions in the N-terminal DBD in the presence of PTSP. We hypothesized that these disordered regions are indicative of partial protein unfolding. The SAXS data revealed that SmcR-apo adopted a radius of gyration (R_g_) of 28.6 Å, whereas SmcR-PTSP adopted a R_g_ of 30.4 Å. R_g_ is defined as the root mean squared deviation of the average of the distance of all electrons from the center of a particle as extrapolated from a Guinier plot of the raw data at low angles. Thus, apo-SmcR adopts a more compact shape compared to SmcR- PTSP, which has a more flexible structure in solution. Consistent with this, the pair-distance distribution functions derived from the scattering profiles, which is a calculation of the distance between any two atoms in a given volume, revealed an apo-SmcR profile with a bell-shaped curve and a single peak of a P(r)/I(0) ratio of 0.00175 at a distance of ∼35 Å, resembling a globular protein with a compact shape. In contrast, the SmcR-PTSP scattering profile showed a smaller peak (P(r)/I(0) ratio of 0.00150) at a distance of ∼35 Å, with an elongated peak that extended to higher r values, suggesting a flexible region of the protein and a potential conformational change in solution (**Fig. 2D, Table S2)**. Overall, these data suggest that SmcR in the presence of PTSP undergoes partial unfolding in solution as compared to SmcR-apo.

### PTSP increases proteolytic degradation of SmcR *in vivo*

From our structural and genetic experiments, we established that PTSP interacts with SmcR by binding in its LBP. Our mutational analyses identified gain-of-function mutants of SmcR that were resistant to PTSP activity *in vivo*. We hypothesized that this resistance phenotype was due to decreased or eliminated PTSP binding. However, this phenotype could also be due to increased expression of SmcR variant proteins compared to wild-type SmcR. Thus, we examined SmcR protein expression in strains expressing wild-type and mutant *smcR* alleles in isogenic *V. vulnificus* mutant strains by western blot analysis. In the wild-type strain, we observed a significant reduction of SmcR protein levels in the presence of PTSP (**Fig. 3A, 3B, Fig. S9A-C**). Conversely, mutant strains expressing SmcR R179C, SmcR H167Q, and SmcR C170F variants did not show altered expression levels of these proteins comparing levels with or without PTSP added (**Fig. 3A, 3B, Fig. S9A-C**). However, the R179C mutant showed overall increased expression compared to wild-type SmcR regardless of the presence or absence of PTSP, correlating with its increased protease activity (**Fig. 3A, 4B, Fig. S9A-C**). We observed similar results with *smcR* mutant alleles when measuring SmcR protein levels in the *E. coli* strains from the genetic screen (**Fig. S9E**).

**Figure 3.**
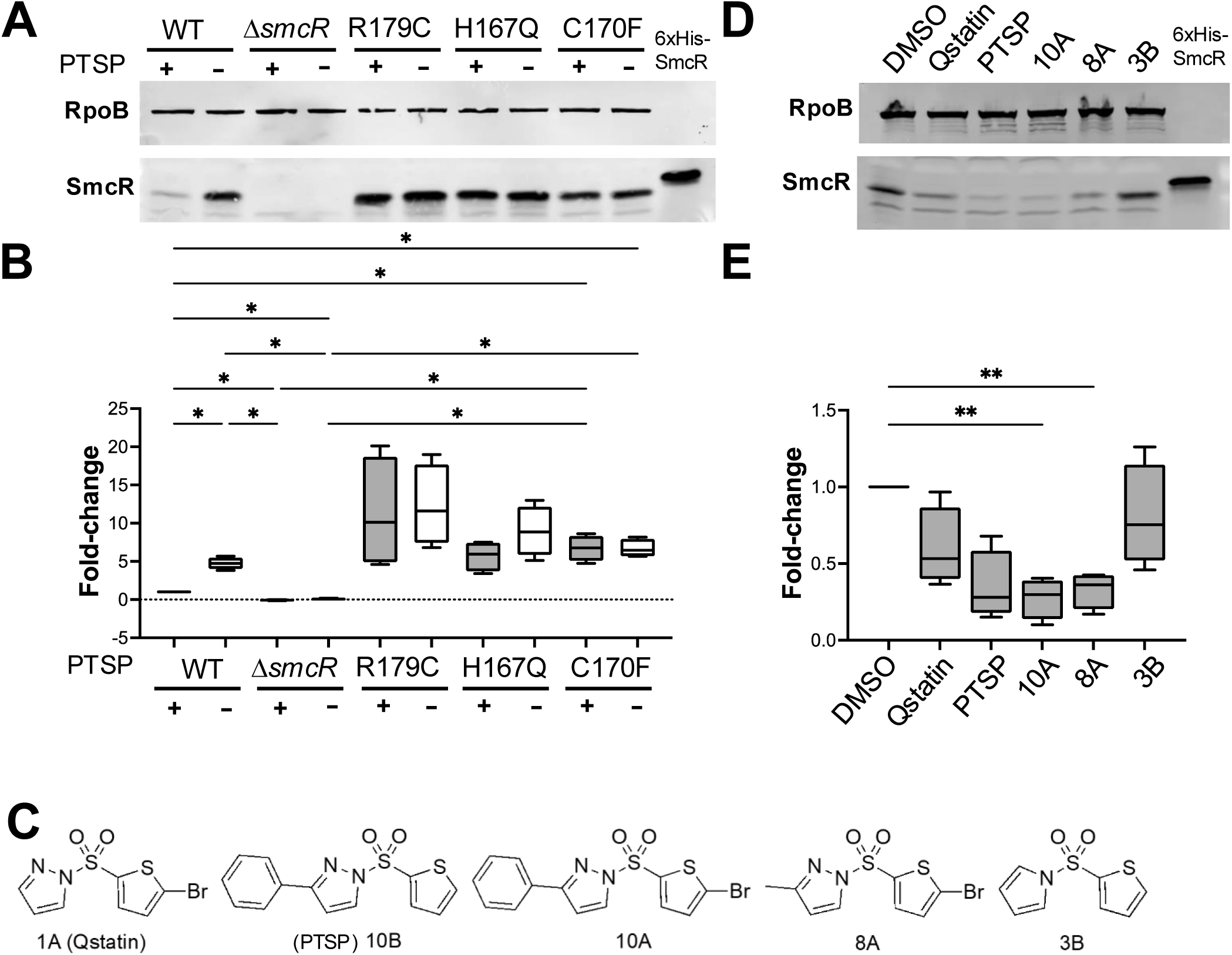
SmcR protein is degraded in the presence of PTSP. (A) Western blot analysis (representative of four biological replicates) of SmcR protein levels in cell lysates from *V. vulnificus* strains wild-type (WT), Δ*smcR smcR* R179C, *smcR* H167Q and *smcR* C170F. Strains were treated with 25 μM PTSP (+) or an equivalent volume of DMSO solvent (-). Anti-RpoB antibodies were used as a loading control. (B) Western blots of four biological replicates (shown in Fig. S9) were quantified with the SmcR band normalized to the RpoB band. Normally distributed data (Shapiro-Wilk test) were analyzed by one-way analysis of variance (ANOVA); *, *p* < 0.05, *n*=4. (C) Structures of thiophenesulfonamides used in panels A, B, D, and E. (D) Western blot analysis (representative of four biological replicates) of SmcR protein levels in cell lysates from WT *V. vulnificus* treated with 25 μM of each thiophenesulfonamide or an equivalent volume of DMSO solvent. Anti-RpoB antibodies were used as a loading control. (E) Western blots of four biological replicates (shown in Fig. S10) were quantified with the SmcR band normalized to the RpoB band. Normally distributed data (Shapiro-Wilk test) were analyzed by ANOVA; **, *p* < 0.01, *n*=4.

**Figure 4.**
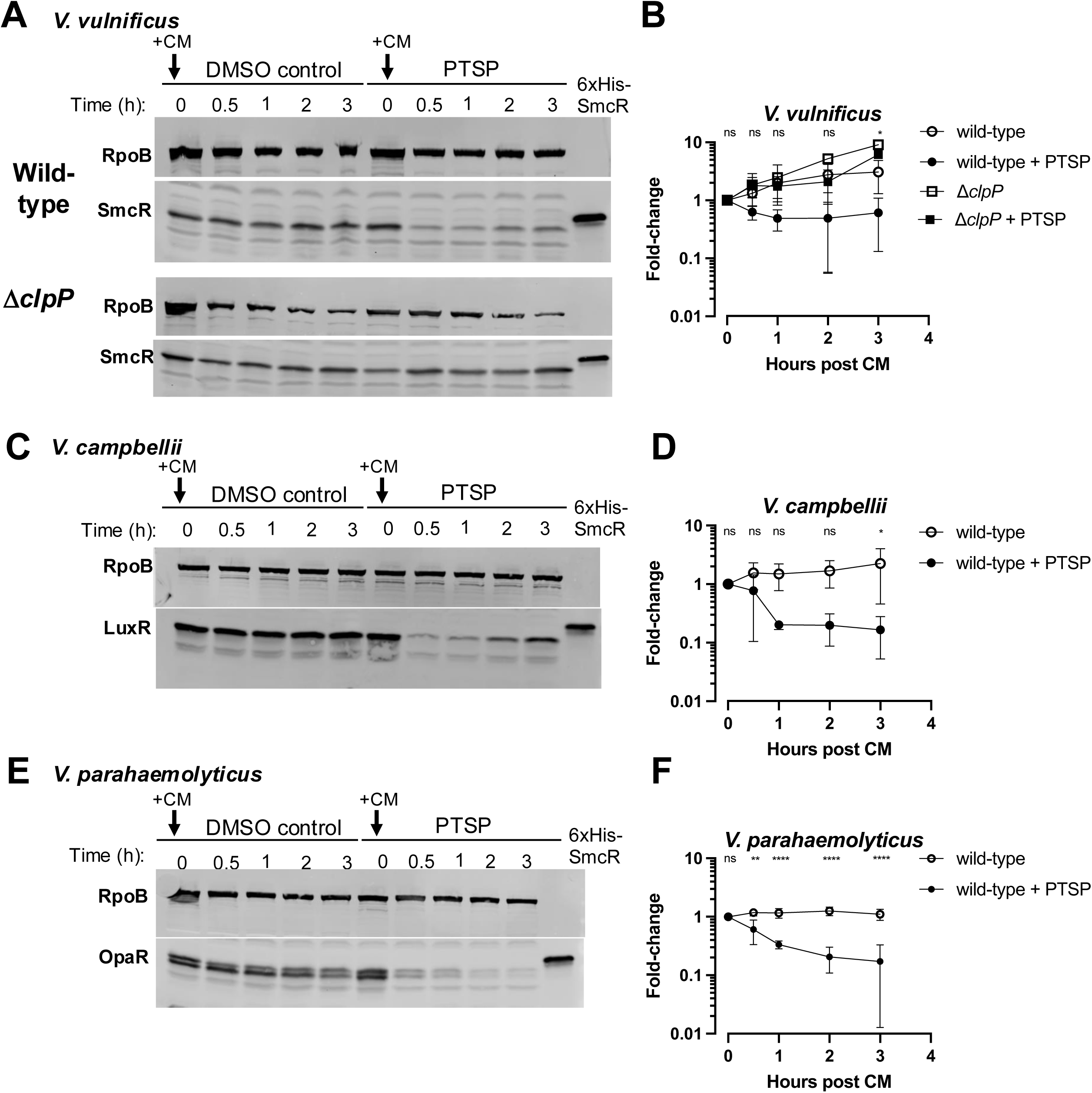
Time-course of SmcR proteins following translation inhibition. (A, C, E) Western blot analysis of cell lysates with anti-LuxR antibodies measuring SmcR from *V. vulnificus* (A), LuxR from *V. campbellii* (C), or OpaR from *V. parahaemolyticus* (E) (all wild-type strains). Cells were either treated with 25 μM PTSP or an equivalent concentration of DMSO solvent and grown to mid-log phase (OD_600_ = 0.5-0.8). Translation was halted by addition of 10 µg/ml of chloramphenicol (CM) at time 0, and pellets were harvested at 0, 30, 60, 120, and 180 mins. Anti-RpoB antibodies were used as a loading control. Each panel is a representative of triplicate biological assays (shown in Fig. S11, S12, S13). (B, D, F) Quantification of SmcR/LuxR/OpaR protein from triplicate biological replicates relative to the RpoB loading control are shown for each *Vibrio* species. Normally distributed data (Shapiro-Wilk test) were analyzed by ANOVA; \**p*<0.05,**, *p*<0.01, ****, *p*<0.0001; *n*=3. For panel B, the significant comparison is between wild-type and wild-type + PTSP.

Next, we tested whether the decrease in SmcR levels upon PTSP addition was specific to this compound or a general effect of thiophenesulfonamide inhibitors. We assayed a select number of thiophenesulfonamide inhibitors that we previously synthesized and characterized *in vivo* (**Fig. 3C**). Each of the compounds tested resulted in an average decrease in SmcR levels compared to the DMSO solvent control (**Fig. 3D, 3E, Fig. S10A-C**). For all compounds, the decreased levels of SmcR correlated with the inhibitor efficacy determined for *V. vulnificus in vivo*. For example, 10A and PTSP were the strongest inhibitors with the lowest IC_50_s against *V. vulnificus* (1 nM and 2 nM, respectively) (43) and also displayed the largest average decreases in SmcR levels *in vivo*. Conversely, Qstatin and 3B had modest-weak efficacy and higher IC_50_ values against *V. vulnificus* SmcR (0.5 µM and 0.9 µM), respectively, and the addition of these Qstatin only moderately-poorly decreased SmcR levels (**Fig. 3D, 3E, Fig. S10A-C**). We conclude that degradation of SmcR correlates with thiophenesulfonamide IC_50_.

To determine the relative stability SmcR in the presence or absence of PTSP, we treated cells with PTSP and monitored SmcR protein stability over time following translation inhibition via chloramphenicol treatment. With this assay, we observed SmcR protein degradation in *V. vulnificus* treated with PTSP as early as 30 minutes post addition of chloramphenicol and lower protein concentrations throughout the time course, whereas cultures not treated with PTSP showed increasing SmcR levels throughout the 3-hour time course (**Fig. 4A, 4B, Fig. S11A-C**).

In our previous publication, we demonstrated that thiophenesulfonamides inhibit the SmcR-type proteins of multiple *Vibrio* species, including *V. campbellii*, *V. parahaemolyticus*, and *V. coralliilyticus*, though to different extents due to the conservation of their LBPs (43). To test if the degradation of SmcR by thiophenesulfonamides is the same mechanism in other species, we assayed degradation in *V. campbellii* and *V. parahaemolyticus* following chloramphenicol treatment. We observed degradation of LuxR (*V. campbellii*) and OpaR (*V. parahaemolyticus*) in cultures treated with PTSP compared to the DMSO controls (**Fig. 4C-F, Fig. S12A-C, Fig. S13A-C**). Much like SmcR, no degradation of LuxR or OpaR was observed in the DMSO-treated cultures, supporting previous observations that these proteins are highly stable *in vivo* (19, 20). From these results, we conclude that treatment of cells with thiophenesulfonamides results in degradation of SmcR/LuxR/OpaR protein levels *in vivo*. Further, we conclude that substitutions of specific PTSP-interacting residues stabilize and/or increase SmcR protein levels *in vivo*.

### PTSP binding stimulates the ClpAP protease to degrade SmcR

SmcR and its homologue HapR have been shown to be degraded by the ClpAP protease *in vivo* (50–52). Further, SmcR is degraded by Lon protease under increased heat stress in *V. vulnificus* clinical isolate MO6-24/0 (50). Thus, we hypothesized that PTSP binding altered SmcR degradation by ClpAP and/or Lon proteases. To test this, we obtained the same strains from these previous studies (a kind gift from Kyung-Jo Lee and Kyu-Ho Lee) and assayed SmcR levels in wild-type, Δ*clpX*, Δ*clpA*, Δ*clpP*, and Δ*lon* mutants of *V. vulnificus* clinical isolate MO6-24/0 in the presence of PTSP or DMSO via western blot. First, we verified that the observations we had made in the *V. vulnificus* ATCC 27562 strain were consistent in the MO6-24/0 strain; we observed SmcR was also degraded in the presence of PTSP (**Fig. 6A**). Next, we assayed the levels of SmcR in isogenic strains containing deletions in *ΔclpA*, *ΔclpX*, *ΔclpP*, and *Δlon* strains. We observed that deletion of *clpA* and *clpP* but not *clpX* or *lon* increased SmcR levels (**Fig. 5A, 5B, Fig. S14**). This result is similar to what was observed for HapR in *V. cholerae* (52). Next, we examined SmcR levels in the PTSP treatment compared to the DMSO control, and again we observed that wild-type SmcR decreased in the presence of PTSP, as well as in the Δ*clpX* and Δ*lon* strains.

**Figure 5.**
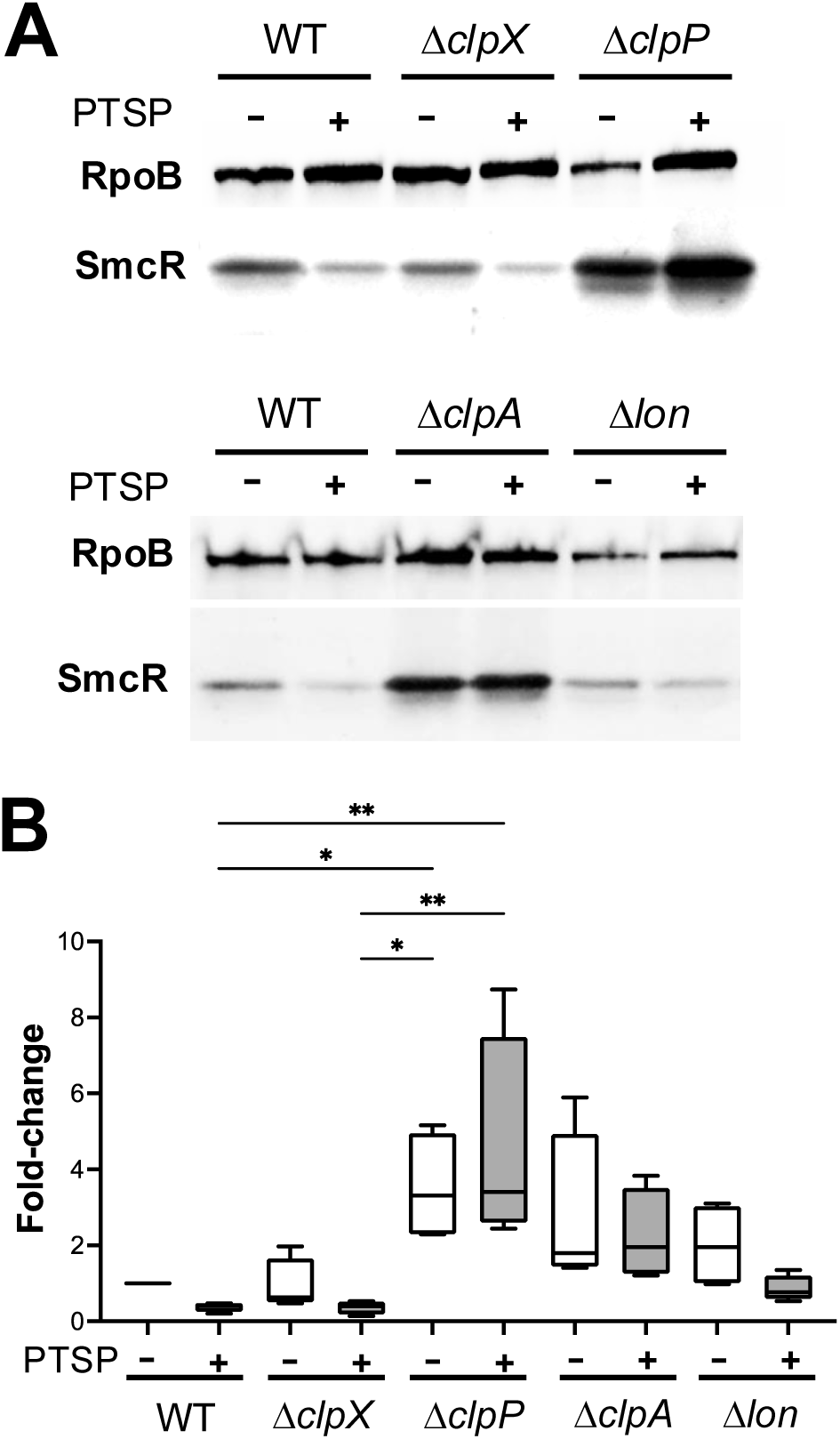
SmcR is proteolyzed by ClpAP. Western blot analysis of cell lysates measuring SmcR from *V. vulnificus* strain MO6-24/0 wild-type (WT), Δ*clpX*, Δ*clpP*, Δ*clpA*, and Δ*lon.* Strains were treated with 25 μM PTSP (+) or an equivalent volume of DMSO solvent (-). Anti-RpoB antibodies were used as a loading control. (B) Western blots of four biological replicates (shown in Fig. S14) were quantified with the SmcR band normalized to the RpoB band. Normally distributed data (Shapiro-Wilk test) were analyzed by ANOVA; \**p*<0.05,**, *p*<0.01, *n*=4.

**Figure 6.**
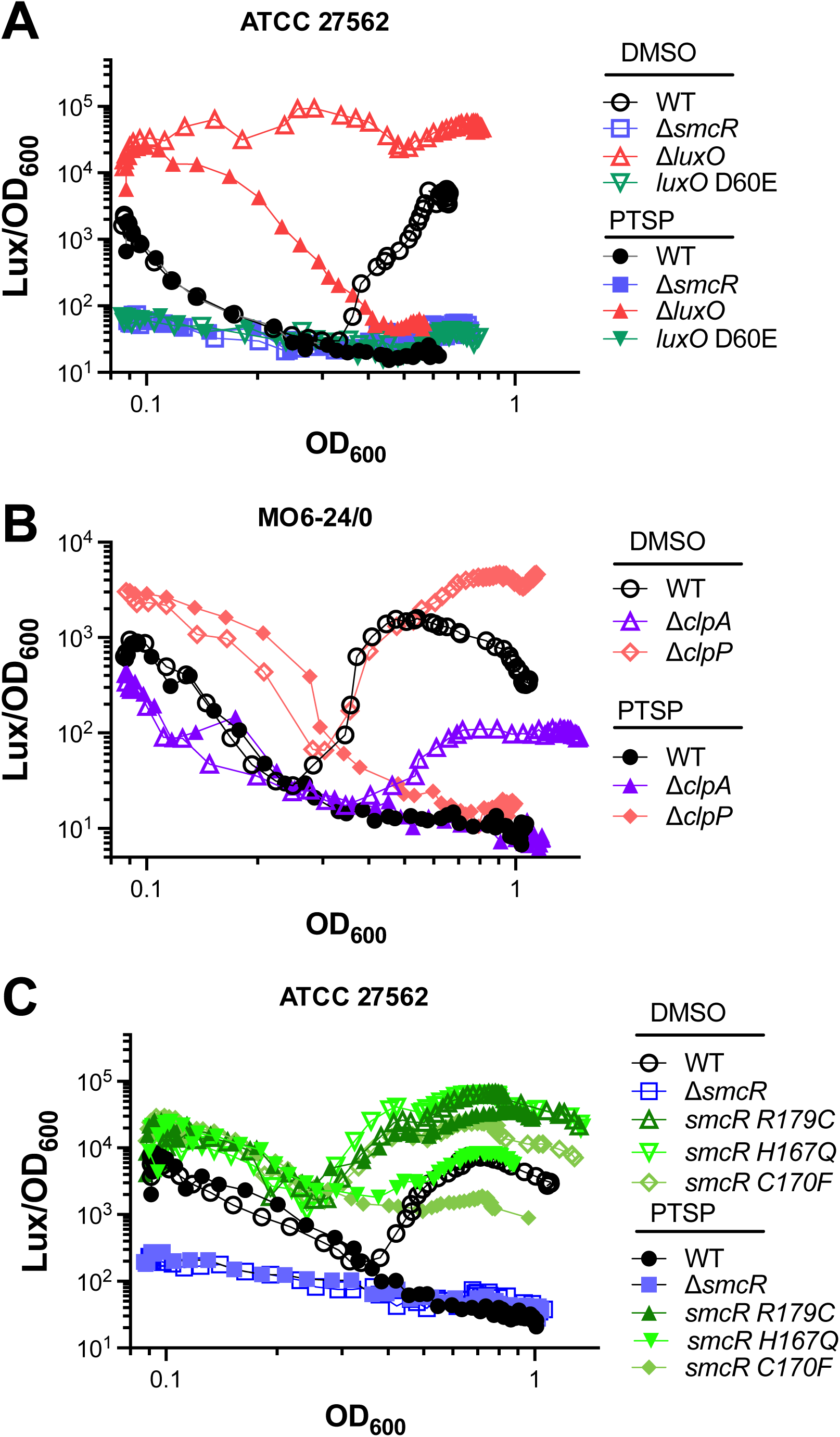
SmcR degradation controls the HCD to LCD transition. (A, B, C) Bioluminescence assays are plotted (Lux/OD_600_) for *V. vulnificus* strains ATCC 27562 (A, C) and MO6-24/0 (B) wild-type (WT) and isogenic mutants. Strains were treated with 25 μM PTSP (+) or an equivalent volume of DMSO solvent (-) Figures are representative of triplicate biological assays (shown in Fig. S15).

However, in the Δ*clpA* and Δ*clpP* strains, SmcR levels were similar with or without PTSP treatment (**Fig. 5A, 5B, Fig. S14**). From these results, we conclude that that SmcR is a substrate for the ClpAP protease, and binding of PTSP to SmcR makes it more susceptible to proteolytic cleavage by ClpAP.

### SmcR degradation controls the timing of the switch in phenotype between high- to low- cell density

The protein levels of SmcR in the cell are controlled by the QS pathway; HCD cultures produce ∼10-fold more SmcR protein than is produced at LCD (19, 53). Because the level of SmcR protein controls the expression of downstream genes, we hypothesized that ClpAP-mediated degradation of SmcR impacts the timing of the switch between densities. To test the effect of SmcR degradation – either by ClpAP or by PTSP – we used bioluminescence expression to examine the QS-state of the cell. We have previously shown that bioluminescence, produced by genes *luxCDABE* and activated by LuxR in *V. campbellii*, is direct readout of SmcR-type protein levels in multiple *Vibrio* species (19, 43, 54, 55). We constructed and tested mutant alleles in *V. vulnificus* that have defined QS phenotypes in other *Vibrio* species. A *luxO D60E* mutant is a phosphomimic that constitutively activates the Qrr sRNAs and behaves as a LCD-locked strain (*i.e*., low expression of SmcR) (**Fig. S1**). Conversely, a Δ*luxO* strain does not activate Qrrs and has constitutively high levels of SmcR, and thus it behaves as a HCD-locked strain (**Fig. S1**). Using a plasmid with the native *luxCDABE* bioluminescence locus derived from *V. campbellii*, we assessed the bioluminescence phenotypes of these mutant strains compared to wild-type. We observed that wild-type *V. vulnificus* produced a standard “U-shaped” curve; as HCD overnight cultures are back-diluted into fresh medium (autoinducers are diluted out), the bioluminescence decreases (**Fig. 6A, Fig. S15, Fig. S16A**). Throughout the growth curve, autoinducers accumulate, and when the cell density reaches quorum, the bioluminescence is activated and becomes maximal at HCD (**Fig. 6A, Fig. S15, Fig. S16A**). Addition of PTSP to wild-type upon back-dilution into fresh medium resulted in complete loss of bioluminescence over time (**Fig. 6A, Fig. S15, Fig. S16A**). The Δ*luxO* strain was constitutively bright in the absence of PTSP, but the bioluminescence decreased to background levels when this strain was treated with PTSP (**Fig. 6A, Fig. S16A**). Both the Δ*smcR* and *luxO D60E* strains were constitutively dark in the presence or absence of PTSP. From these data, we conclude that addition of PTSP to wild-type or Δ*luxO* cultures results in loss of bioluminescence, indicating that SmcR level and degradation by PTSP is epistatic to the Δ*luxO* allele.

Using this bioluminescence assay, we sought to examine the impact of ClpAP degradation of SmcR during the transitions between HCD to LCD, and also from LCD to HCD. We hypothesized that deletion of either *clpA* or *clpP* would increase SmcR levels and thus bioluminescence expression throughout the curve, delating the transition between HCD to LCD and increasing the transition between LCD to HCD. First, comparing only the untreated strains (DMSO control), the Δ*clpP* mutant was delayed in the HCD to LCD transition compared to wild-type and increased to a higher final bioluminescence level (**Fig. 6B, Fig. S16B**). Conversely, the Δ*clpA* mutant had lower bioluminescence levels compared to wild-type throughout the curve (**Fig. 6B, Fig. S16B**). All PTSP-treated strains decreased levels of bioluminescence that reached basal levels at the end of the curve (**Fig. 6B, Fig. S16B**). Initially, we were puzzled by the Δ*clpA* result; our hypothesis had been correct for Δ*clpP*, but not Δ*clpA*. However, upon reading literature in the field, previous studies have shown that in *Caulobacter*, deletion of *clpA* increased Lon levels (56). Thus, we hypothesize that in the absence of *clpA*, Lon targets SmcR for degradation more than in wild-type. Further, we note that treatment of the Δ*clpP* mutant with PTSP still resulted in loss of bioluminescence. This may also be due to increased levels of Lon activity. We note that the Δ*clpA* and Δ*clpP* strains are in the MO6-24/0 *V. vulnificus* background, whereas the other strains in this experiment were in ATCC 27562. To verify that the phenotypes we observed also occurred in the ATCC 27562 strain, we constructed isogenic Δ*clpA* and Δ*clpP* mutants in that strain background and assayed bioluminescence. The results were similar to the MO6-24/0 strain: in the absence of PTSP treatment, the *ΔclpA* strain had slightly decreased bioluminescence compared to wild-type, whereas Δ*clpP* had increased bioluminescence (**Fig. S17A-C**). After treatment with PTSP, both mutants had basal levels of bioluminescence, indicating that the SmcR protein in these mutant strains is still susceptible to PTSP- mediated degradation.

To further determine the role of SmcR degradation in HCD-LCD-HCD transitions, we assessed the three mutant *smcR* alleles R179C, H167Q, and C170F, that increased SmcR stability to varying degrees. We observed several distinctions in transition timing in untreated cultures (**Fig. 6C, Fig. S16C**). First, each of the mutants exhibited a consistent delay in the switch from HCD to LCD compared to wild-type, with the R179C mutant being the strongest phenotype. Second, each of the mutants exhibited strong differences in the transition from LCD to HCD, both earlier in timing and more bioluminescence production than wild-type, again with R179C being the strongest phenotype. Third, all three mutants also exhibited higher overall levels of bioluminescence than the wild-type throughout the growth curve. These data are consistent with our biochemical assays and western blot analyses (**Fig. 1**, **Fig. 2**, **Fig. 3A)**. Next, we examined the effects of adding PTSP to the cultures at back-dilution into fresh medium. We observed that there was a decrease in the final bioluminescence levels for *smcR* H167Q and *smcR* C170F compared to untreated cultures, indicating that these two mutants were still somewhat sensitive to PTSP treatment (**Fig. 6C, Fig. S16C**). However, the R179C mutant produced higher levels of bioluminescence than wild-type at HCD even when treated with PTSP (**Fig. 6C, Fig. S16C**). Collectively, from these results, we made three conclusions: 1) ClpAP turnover of SmcR affects both QS transition states, 2) when cultures are treated with PTSP, in the absence of ClpAP, another mechanism of SmcR turnover occurs, and 3) the SmcR variant R179C is resistant to protein degradation.

## Discussion

*Vibrio* species have served as cornerstone models for studying numerous bacterial behaviors, including biofilm formation, virulence factor secretion, host-pathogen interactions, bacteriophage-host evolution, signaling, and more (57–65). QS was discovered in *Vibrio* species and much of the system knowledge in the field was established through research on QS in *Vibrio cholerae, Vibrio fischeri,* and *Vibrio campbellii* (in a strain previously classified as *V. harveyi*). The identification of the master transcriptional regulators at the center of the QS circuit that we call LuxR/HapR proteins led to the discovery of key gene families regulated by QS: biofilms, motility, competence, toxins, etc. The first structural assessment of this *Vibrio*-specific protein family focused on HapR, followed by similar studies of *V. vulnificus* SmcR and *V. alginolyticus* LuxR. The high amino acid identity of these proteins is even surpassed by the nearly identical tertiary structures of these proteins, which unsurprisingly bind to nearly identical DNA sequences *in vivo*. The literature is thorough in examinations of LuxR/HapR promoter DNA binding, gene regulons, and protein-protein interactions that drive the global activity of these proteins (18, 66–68). However, one key attribute of these proteins had not been evaluated: the ligand binding potential. As a subset of the massive TetR protein family, LuxR/HapR proteins have a LBP clearly defined by an open channel lined with hydrophilic residues and a hydrophobic core (26). Indeed, in every X-ray crystal structure of the “apo” protein, there is density observed in the LBP, even though these proteins are purified from *E. coli* (25, 26, 43). Yet no phenotype has been determined for one of these proteins without its native ligand. Rather, a few studies have shown that inhibitors can bind LuxR/HapR proteins, decreasing their transcriptional regulation activity *in vivo*. Studies of thiophenesulfonamide inhibitors such as PTSP and Qstatin have likewise shown that these molecules bind specifically to LuxR/HapR proteins with varying affinities (25, 43). However, the mechanism of action was not known. Assays *in vitro* for DNA binding yielded no significant differences in binding affinity, yet *in vivo* the addition of these compounds clearly blocked LuxR/HapR activity.

Here, we identified the mechanism of thiophenesulfonamide inhibition of LuxR/HapR proteins via ClpAP degradation. We propose a model for SmcR degradation by ClpAP and its impact on QS transition states in **Figure 7**. Our data show that SmcR binds thiophenesulfonamides in its LBP, and addition of these compounds *in vitro* alters SmcR protein folding dynamics. We determined that thiophenesulfonamides such as PTSP drive SmcR proteolysis via the ClpAP protease, hence blocking its activity *in vivo*. Thus, we have uncovered the reason that *in vitro* DNA binding assays were not altered by addition of PTSP nor Qstatin but only *in vivo* activity was changed. Armed with this clear and measurable proteolysis phenotype driven by inhibitor binding, we now aim to uncover the role of the native ligand in LuxR/HapR function *in vivo*.

**Figure 7.**
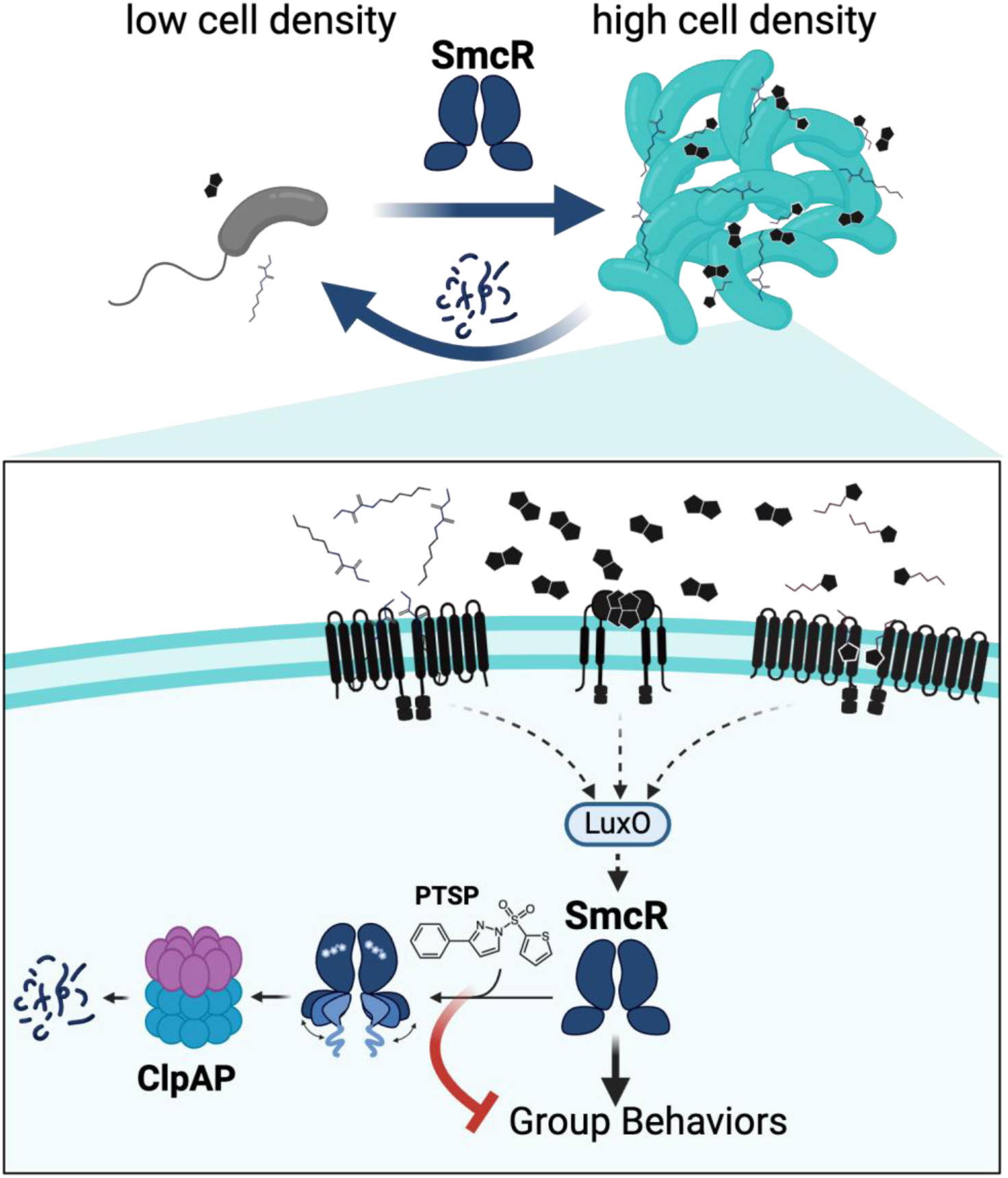
Model of the role of SmcR degradation in quorum sensing transitions. At low cell densities, low levels of autoinducers result in low levels of SmcR. As cells transition to high cell density, more SmcR is produced, which drives expression of group behavior genes. Our model is that SmcR is bound to its native ligand (unknown) and is most active at high cell density. As cells transition back to low cell density, degradation of SmcR by ClpAP increases the speed of this transition. Addition of inhibitor thiophenesulfonamide PTSP to cultures displaces the native ligand, driving an allosteric conformational change in the DNA binding domains of each SmcR monomer. We propose that the unfolded N-terminus is recognized by ClpAP and degraded, which blocks expression of group behavior genes. Image created with bioRender.

The next question we aim to focus on is: how does inhibitor binding to SmcR result in increased proteolysis? We hypothesize that this is accomplished via conformational changes, as is often observed with ligand binding for TetR proteins (69). Although we have successfully solved the structure of SmcR bound to PTSP, the structure itself does not indicate large conformational changes that might explain how SmcR is targeted for proteolysis when bound to PTSP. This may be due to the limited nature of X-ray crystallography in that the structures are not determined in solution, and not all possible conformations may form crystals. Based on our SAXS profiles and pair-distance distribution functions, SmcR bound to PTSP in solution adopts a more elongated and flexible conformation. This may be a potential mechanism for targeted proteolysis. We thus propose a model in which PTSP (or thiophenesulfonamides in general) binds to the LBP of each monomer in the LuxR/HapR protein, resulting in a conformational change that increases proteolysis of the dimer by the ClpAP protease (**Fig. 7**), possibly by exposing a ClpA- dependent degron. We also aim to examine whether there is an adaptor protein such as ClpS required for proteolysis by ClpAP. Notably, the *clpS* mutant did not exhibit a change in HapR protein levels in *V. cholerae* (52), suggesting that ClpS is not required for ClpAP-mediated proteolysis in that system. We also note that even in a Δ*clpP* strain, PTSP addition resulted in loss of bioluminescence, suggesting that another protease may contribute to SmcR degradation. We hypothesize that the other protease activity is from Lon. Several lines of evidence support this: 1) deletion of *clpA* in *Caulobacter* increases Lon protein levels (56), 2) Lon has been shown to target SmcR during heat stress (46), and 3) degradation-resistant alleles of *smcR* had increased bioluminescence even in the presence of PTSP, indicating that these alleles are resistant to all protein degradation. Future genetic studies of the genes required for LuxR/HapR protein degradation and biochemical studies of ligand binding will inform the mechanism of thiophenesulfonamide-driven protein proteolysis.

Recent studies have shown that QS inhibitors are feasible alternatives to traditional antibiotics with demonstrated efficacy in infection models (31, 32). As these inhibitors do not affect bacterial viability, they are hypothesized to generate less selective pressure for evolving resistance (33, 34). These factors make them attractive targets for *Vibrio* QS inhibition and possibly treatments for *Vibrio* diseases. For example, Qstatin markedly decreased virulence of three *Vibrio* species against the brine shrimp *A. franciscana* (29). Together, these results provide insights into the mechanism of action of an effective QS inhibitor that may be useful in controlling *Vibrio* virulence in aquaculture.

Broadly, our discovery that ClpAP-mediated degradation impacts *V. vulnificus* QS phenotype states has many implications that are relevant to cell survival and proliferation in native environments. Our current model is that the levels of SmcR are controlled by ligand binding, whether the ligand is an agonist or antagonist. The turnover of SmcR by ClpAP protease (and likely also another protease such as Lon) mediates transitions between cell density states. We can envision different scenarios for both transitions. When cells are at HCD and shifted to LCD, such as dispersal from a biofilm or host, QS signal concentrations would quickly drop. However, the level of SmcR starting at its most concentrated would not respond as quickly as the kinase/phosphorylation state of the receptors and regulators or the mRNA turnover. Thus, the proteolytic turnover of SmcR would increase the rapidity with which cells could alter gene expression to be in the LCD state. When cells transition from LCD to HCD, as cells colonize a host and build a biofilm, the sensitivity of the transition from LCD to HCD is dictated by the accumulation of SmcR protein. The ClpAP-proficient wild-type strain exhibited a longer time to reach “quorum”, suggesting that the turnover of SmcR controls the timing of LCD to HCD transition. Further exploration of the mechanisms of ClpAP-mediated SmcR proteolysis and how this is impacted by putative native ligands will be necessary to fully understand the impact of SmcR turnover on QS gene expression timing.

## Materials and Methods

### Bacterial strains and media

All strains and plasmids from this study are listed in Table S3 and Table S5. *Escherichia coli* strains DH10B and S17-1λpir were used for cloning, and BL21(DE3) was used for the overexpression of SmcR proteins. All *E. coli* strains and derivatives were grown at 37°C shaking at 275 rpm in Lysogeny Broth (LB) medium with appropriate antibiotics. *V. vulnificus* and *V. campbellii* strains and derivatives were grown shaking at 275 rpm at 30°C in LB Marine (LM) medium (LB with 2% NaCl final concentration) with appropriate antibiotics. Antibiotics were used at the following concentrations: kanamycin 40 μg/ml or 100 μg/ml (*E. coli* or *Vibrio*, respectively), chloramphenicol (10 μg/ml), and tetracycline (10 μg/ml).

### Compound synthesis

Qstatin, PTSP, and all other derivatives used in this study were either synthesized as described in Newman *et. al*, 2021.

### Random mutagenesis and flow cytometry

The *smcR* mutant alleles were created using GeneMorph II EZClone Domain Mutagenesis Kit (Stratagene) as per the company protocol targeting the *smcR* gene encoded on plasmid pJN22, which expresses *V. vulnificus smcR* under control of the IPTG-inducible P*_tac_* promoter (43). Mutant libraries were transformed into S17-1λpir competent cells, pooled, and stored at -80°C. DNA was extracted from the library and then transformed into competent cells of *E. coli* strain DH10B containing the pJV064 reporter plasmid, selected on LB plates with kanamycin and chloramphenicol, pooled, and stored at -80°C. Overnight cultures of the pooled library were back-diluted 1:1,000 into fresh LB media with chloramphenicol (10 µg/ml), kanamycin (40 µg/ml) and IPTG (50 μm), treated with either DMSO or PTSP (25 μm) and grown shaking at 30°C overnight. Aliquots were prepared for flow cytometry by making 1:100 or 1:1,000 dilutions in PBS 30 mins prior to flow. Mutant strains resistant to PTSP were isolated by sorting cells with high GFP and low mCherry fluorescence on a FACS Aria II. Following sorting, colonies were inoculated individually into fresh medium, grown to stationary phase, and stored at -80°C.

#### Dual-reporter E. coli bioassay

Overnight *E. coli* cultures containing either pJN22 or empty vector control pMMB67EH-kanR and the pJV064 reporter were diluted 1:1000 into LB with chloramphenicol (10 µg/ml) and kanamycin (40 µg/ml) and aliquoted into black-welled, clear-bottomed 96-well plates to a final volume of 150 µl. PTSP was resuspended in DMSO at a stock concentration of 10 mM and added to *E. coli* cultures at a final concentration of 25 µM unless otherwise stated. DMSO was added as a negative control at equal final concentration. 50 µM IPTG was also added to the wells to activate expression of *smcR*. The 96-well plates were covered in microporous sealing tape and grown for 16-18 hr shaking at 275 rpm at 30°C. The OD_600_ and GFP and mCherry fluorescence were measured on a BioTek Cytation plate reader.

### Molecular methods

All PCR reactions were performed using Phusion HF polymerase (New England Biolabs). All oligonucleotides were purchased from Integrated DNA Technologies (IDT), and those used in this study are listed (Table S5). All mutant strains (Table S3) were confirmed by DNA sequencing (Eurofins). Three plasmids were constructed by Twist Biosciences.

### SmcR protein purification

His-tagged wild-type and mutant SmcR proteins were expressed and purified as described previously (70) using an Akta Pure L FPLC with His-Trap and gel-filtration columns. For induction of His-tagged wild-type and mutant His-SmcR in presence of PTSP, induction media were supplemented with 50 μM of PTSP dissolved in DMSO. No additional PTSP was added to buffers during purification. Proteins were stored in gel filtration buffer (25 mM Tris, 300 mM NaCl, 1 mM DTT, 5 mM EDTA at pH 7.5) at 4°C.

#### *In vivo* protein turnover assay

*Vibrio* strains were streaked on LM plates with kanamycin (100 µg/ml), spectinomycin (200 µg/ml), or trimethoprim (10 µg/ml) and incubated overnight at 30°C. Colonies were inoculated in 2 ml LM with kanamycin (100 µg/ml), spectinomycin (200 µg/ml), or trimethoprim (10 µg/ml) and incubated at 37°C until an OD_600_ = ∼1.0. Cultures were diluted in 25 ml of LM to an OD_600_=0.05 and incubated at 30°C with shaking (275 rpm) until ∼OD_600_ = 0.5 to 0.8. Chloramphenicol (10 ug/ml) was added to the cultures and cells were harvested at different time points thereafter (0, 30, 60, 120, and 180 minutes) by centrifuging at 14,000 rpm for 2 minutes. Pellets were resuspended in lysis buffer (1 ml 1x Bugbuster, 50 µg/ml lysozyme,1µl Benzonase, 1X Protease Inhibitor) to 0.01 ODs/µl to stop the reactions. Processed samples were analyzed by western blot as per previous protocol (17) using anti-LuxR antibodies, and anti-RpoB antibodies were used as loading control.

### V. vulnificus strain construction

Isogenic strains containing mutations in *smcR* were generated by mutating wild-type *V. vulnificus* strain ATCC 27562. All isogenic mutant strains were generated using natural transformation via *tfoX* induction and MuGENT (71, 72) via oligonucleotides listed in Table S4. First, derivative cas-vv006 was constructed in which a trimethoprim-resistance (Tm^R^) cassette was inserted downstream of *smcR* at the native locus. The cas-vv006 strain was then used as a template to amplify *smcR*-Tm^R^ and generate targeting DNA to introduce substitutions into *smcR* via natural transformation. These DNA products were then introduced into wild-type *V. vulnificus* and selected for on LM plates with trimethoprim (10 µg/ml). All isogenic mutant strains were confirmed by sequencing at the *smcR* locus.

### V. vulnificus growth curves

*Vibrio* strains were inoculated in 5 ml LM or M9 medium (supplemented with casamino acids and 1% tryptone) overnight at 30°C shaking at 275 rpm with kanamycin (100 µg/ml), gentamicin (10 µg/ml), spectinomycin (200 µg/ml), or trimethoprim (10 µg/ml) to select for reporter plasmids as needed.

Overnight cultures were back-diluted 1:100,000 in LM or M9 medium with antibiotics, treated with 25 μm PTSP or an equal volume of DMSO and transferred into clear-bottomed 96-well plate. The plate was covered and incubated in a BioTek Cytation plate reader overnight with shaking at 275 rpm at 37°C for 22 hr. OD_600_ was measured at 30 min intervals throughout the overnight incubation period.

### Protease assays

*Vibrio* strains were inoculated in 5 ml LM with kanamycin (100 µg/ml), spectinomycin (200 µg/ml), or trimethoprim (10 µg/ml) and incubated overnight at 30°C shaking at 275 rpm. Overnight cultures were back diluted 1:1,000 in 10 ml LM with kanamycin (100 µg/ml), spectinomycin (200 µg/ml) or trimethoprim (10 µg/ml) and the cell mixture was equally aliquoted into test and control sets, with 5 ml culture per set. PTSP was added to test reactions at a concentration of 25 µM while DMSO was added as a negative control at equal final concentrations into control reactions. These cultures were grown for 16-18 hrs shaking at 275 rpm at 30°C. After incubation, OD_600_ was measured (1:10 dilution of overnights), and 1 ml of overnight undiluted cultures were pelleted in 1.7 ml micro centrifuge tube at 13,000 rpm for 10 min at room temperature. 100 µl supernatant was transferred to a new clear 1.7 ml microcentrifuge tube and incubated with 400 µL of 1% azocasein (0.01 mg/ml dissolved in dH_2_O) for 2 hrs 30 min at 37°C. The reaction was stopped by addition of 600 µl 10% trichloroacetic acid, and the tubes were further incubated on ice for 30 min, then centrifuged at 13,000 rpm for 5 min at room temperature. 800 µl of the protease reaction was transferred to a new clear microcentrifuge tube, and 200 µl of 1.8 N NaOH was added.

OD_420_ was measured in a desktop spectrophotometer. Protease activity was calculated by dividing OD_420_ by OD_600_. Each assay was performed in technical and biological triplicates.

### Western blots

*Vibrio* strains were inoculated in 5 ml LM with kanamycin (100 µg/ml), spectinomycin (200 µg/ml), or trimethoprim (10 µg/ml) and incubated overnight at 30°C shaking at 275 rpm. Overnight cultures were back diluted 1:1,000 in 10 ml LM with kanamycin (100 µg/ml), spectinomycin (200 µg/ml), or trimethoprim (10 µg/ml) and the cell mixture was equally aliquoted into test and control sets, with 5 ml culture per set. PTSP was added to test reactions at a concentration of 25 µM while DMSO was added as a negative control at equal final concentrations into control reactions. These were incubated at 30°C shaking at 275 rpm. Cells were harvested after 16 hours growth overnight otherwise specified and resuspended in lysis buffer (1 ml 1x Bugbuster, 50µg/ml lysozyme,1µl Benzonase and 1X protease inhibitor) to an OD_600_ of 0.01/µl and incubated at room temperature for 10 minutes. SDS sample buffer was added to 10 µl lysate with fresh β-mercaptoethanol (10%), and samples were boiled for 10 mins at 95°C. Western blot analyses were performed as previously described (19), except that new anti-LuxR antibodies were generated through Cocalico against purified LuxR protein. In addition, the western blots were probed with anti-RpoB as a loading control as previously described (Neoclone) (73).

### Purification of SmcR:PTSP for crystallographic studies

His-tagged wild-type SmcR-PTSP was expressed and purified via Ni-affinity chromatography as previously described. Induction of expression was performed in induction media supplemented with 50 µM PTSP in DMSO. Clarified cell lysates were incubated with Ni-NTA running buffer equilibrated NEBExpress Ni Resin for 1 hour with rocking at 4°C. Incubated resin was applied to a gravity column and washed with 100 mL Ni-NTA running buffer. His-tagged wild-type SmcR-PTSP was eluted in 10 mL Ni-NTA elution buffer in 1 mL fractions. Peak fractions, confirmed by SDS-PAGE, were pooled, and concentrated to 1 mL over a 10 kDa MWCO filter (Amicon) for injection on a Superdex-200 size exclusion column (Cytiva) equilibrated with gel-filtration buffer (25 mM Tris-HCl pH 7.5, 200 mM NaCl, 1 mM DTT). Peak fractions, determined by 280 nm absorbance and confirmed by SDS-PAGE, were pooled and concentrated to 5.2 mg/mL in gel-filtration buffer. SmcR:PTSP was stored at 4°C for immediate crystallographic screening.

### Isothermal calorimetry analysis

PTSP was re-suspended in 100% DMSO to a final concentration of 333 mM and serial diluted into a solution containing decreasing amount of DMSO and increasing amount of gel filtration buffer (200 mM NaCl, 25 mM Tris pH 7.5) to a final concentration of 10 μM PTSP with 0.15% DMSO in the buffer. SmcR-wild-type and SmcR-R179C proteins were purified as described. Purified proteins were then denatured through dialysis at room temperature in a buffer containing 500 mM NaCl, 25 mM Tris pH 7.5, 1 mM DTT and 8 M Urea for 4 h, then switched to buffer containing 500 mM NaCl, 25 mM Tris pH 7.5, and 1 mM DTT for 4 h to remove any potential natively bound ligands. Aggregates were also removed. The re-natured proteins were then dialyzed against ITC buffer (200 mM NaCl, 25 mM Tris pH 7.5 and 0.15% DMSO) overnight, concentrated to ∼4.5-6 mg/mL. A MicroCal PEAQ-ITC (Malvern) was used to measure protein-ligand binding affinity at 25°C. 84μM to 120 μM proteins (4.4 mg/mL to 6 mg/mL, calculated as dimers) in the syringe were titrated into 10 μM PTSP in the cell, with 500 rpm stirring speed and thermal power recorded every 10s. Data were fitted and analyzed through the PEAQ-ITC Analysis software (Malvern).

### Crystallographic studies

Previous crystallographic conditions for the determination of the SmcR-apo and SmcR-Qstatin structures did not yield crystals for SmcR-PTSP. Purified SmcR-PTSP (5.2 mg/mL) in gel-filtration buffer was screened using a variety of kit solutions (Nextal and Hampton) using a Gryphon Crystallization Robot (ART Robbins Instruments) and incubated at RT for 4 days. Optimization screens were performed using the hanging drop vapor diffusion method in 24-well crystallization trays. Initial SmcR-PTSP crystals were grown by mixing 1 µL protein solution with 1 µL well solution (2.0 M MgSO_4_, 0.1 M MES pH 6.5) and sealed over a 1 mL well solution. After incubation for 24 hours at RT, SmcR-PTSP crystals were harvested, transferred to a PCR tube containing well solution, and vortexed. Fragmented crystals were used to streak-seed new drops of 1:1 protein:well solution to optimize crystal order and size. We thank Ted and Artemis Andrus-Paczkowski for the generous donation of their shed whiskers. Seeded SmcR-PTSP crystals formed after 24 hours incubating at RT, were batch harvested and soaked in cryoprotectant (20% v/v glycerol, 0.1 M MES pH 6.5, 1.6 M MgSO_4_) for 10 min. After the cryoprotecting step, single crystals were mounted on a MiTeGen loop and flash frozen and stored under liquid nitrogen.

### Data collection and structure solution

X-ray diffraction data were collected at Brookhaven National Laboratory National Synchrotron Light Source-II on the NYX beamline. Diffraction images were processed in XDS and data reduction was performed using CCP4-8.0.013 and Aimless (74). Automated and manual structure solutions were generated via molecular replacement using *MrBUMP* and *SHELXE-MR*, respectively (75). Molecular replacement in MrBUMP fit two copies of *V. vulnificus* SmcR-Qstatin (PDB: 5X3R, seq. identity 100%) chain A with waters, sulfate ions, and Qstatin removed. Results from MrBUMP were used for molecular replacement in SHELXE-MR. Alternating rounds of manual and automated refinement were performed using Coot v0.9.8.8 and REFMAC5 (76). Structural validation was performed using Molprobity. Graphics were created using Chimera (77).

### Small angle X-ray scattering

SAXS data were collected at the Cornell High Energy Synchrotron Source (CHESS) 7A1 beamline over multiple runs. For all runs, a 250 µM x 250 µM X-ray beam with a wavelength of 1.094 Å at 7.1x10^11^ photons s^-1^ was used and centered on a quartz capillary sample cell with a path length of 1.5 mm. 10, 1-second images were collected using a dual Pilatus 100K-S detector system (Dectris, Baden, Sweden), covering a range of *q* ≈ 0.0079-0.45 Å^-1^. The 1D SAXS profiles were grouped by sample and averaged, followed by buffer subtraction (25 mM Tris-HCl pH 7.5, 200 mM NaCl, and 1 mM DTT) within BioXTAS RAW (v2.2.1) (78). The radius of gyration (*R_g_*) and forward scattering intensity *I*(0) were estimated by Guinier fit analysis, and the pair distance distribution function analysis was done in GNOM (79) utilizing the ATSAS software suite (80)v3.2.1-1) implemented in BioXTAS RAW (Hopkins et al., 2017) (v2.2.1). The molecular weights were estimated in a concentration independent manner from the Porod volume, *V*_p_ (81). Single-state modelling was performed using the FoXS server (https://modbase.compbio.ucsf.edu/foxs/)(82, 83).

### Bioluminescence curves

Bioluminescence assays were performed in *V. vulnificus* ATCC 27562 and MO6-24/0 strains containing IPTG-inducible pCS38 *luxCDABE* reporter plasmid (70). Overnight cultures of *V. vulnificus* were diluted to 1:100000 in M9 minimal medium (supplemented with casamino acids and 1% tryptone) with gentamycin (100 µg/ml) and treated with 25 uM PTSP or an equal volume of DMSO in 96-well clear bottom black assay plates. The plates were incubated shaking at 30°C for 48 h in the BioTek Cytation 3 Plate Reader or BioTek Synergy H1 Plate Reader with bioluminescence and OD_600_ readings taken every 30 minutes.

## Data analysis

All kinetic data were processed using GraphPad Prism software (GraphPad Software, Inc). For statistical significance tests, data were analyzed using GraphPad Prism to perform tests to conform the normal distribution of the data (Shapiro-Wilk test) with subsequent parametric tests for significance (ANOVA and/or t-tests) as indicated in the figure legends.

## Data availability

The X-ray structure data for SmcR-PTSP has been deposited (PDB: 8W39). All other data that support the conclusions of this study are contained or described within the text or supporting information.

## Acknowledgments

We thank Dr. Giovanni Gonzalez-Gutierrez and Juan Nysschen at Indiana University for technical support, and Dr. Kyung-Jo Lee for sending us *V. vulnificus* MO6-24/0 Δ*clpA*, Δ*clpP*, Δ*clpX*, and Δ*lon* mutant strains. We also thank Cristina Landeta and the Landeta lab for helping with western blot protocols. We thank Dr. Peter Chien for helpful discussions about ClpAP and Lon. This research used resources of the National Synchrotron Light Source II (NSLS-II), a U.S. Department of Energy (DOE) Office of Science User Facility operated for the DOE Office of Science by Brookhaven National Laboratory under Contract NO. DE-SC0012704. The authors thank Kevin Battaile, Director of the NYX beamline at NSLS-II at Brookhaven National Laboratory, for technical assistance during data collection. We would like to thank the members of van Kessel lab for their support and thoughtful discussions. Cartoon diagrams in Figures S1 and 7 were created with bioRender.

## Funding and additional information

The project was supported by NIH training grant T32GM132066 and NSF REU award DBI-175710 to MWB, NIH grant R01GM14436101 to JEP, and NIH grant R35GM124698 to JVK. This work is based on research conducted at the Center for High-Energy X-ray Sciences (CHEXS), which is supported by the National Science Foundation (BIO, ENG and MPS Directorates) under award DMR-1829070, and the Macromolecular Diffraction at CHESS (MacCHESS) facility, which is supported by award 1-P30-GM124166-01A1 from the National Institute of General Medical Sciences, National Institutes of Health, and by New York State’s Empire State Development Corporation (NYSTAR).

## Author contributions

JVK, JEP, LCB, and TAR designed the research, TAR, CPM, MWB, BL, CAS, ADP, FJA, JEP, and JVK performed experiments, JVK, JEP, LCB, CPM, and TAR analyzed data, and JVK, JEP, LCB, TAR, and CPM wrote the manuscript.

