## Supplemental Information for "Protease-mediated degradation of the master transcription factor controls quorum sensing-state transitions in *Vibrio*"

Supplementary Information

Video S1

Tables S1-S5

Figures S1-S16

**Video S1.** X-ray crystal structure of SmcR bound to PTSP (PDB: 8W39).

**Table S1.** Crystallographic statistics. Values in parentheses represent the highest resolution bin.

| | |
|---|---|
| Space group | P21 21 21 |
| Unit cell dimensions (Å) | 58.01, 64.65, 138.73, 90, 90, 90 |
| Resolution (Å) | 47.29-2.10 (2.10-2.18) |
| Total reflections | 208777 (17499) |
| Unique reflections | 31260 (2538) |
| Completeness (%) | 99.3 (99.0) |
| $R_{merge}$ | 0.125 (0.217) |
| Clash score | 6.51 |
| <I>/<sigma> | 15.1 (7.0) |
| Total atoms | 1409 |
| $R_{work}$ (%) | 26.1 (36.3) |
| $R_{free}$ (%) | 29.1 (36.0) |
| Wilson B-factor | 22.0 |
| **Number of non-hydrogen atoms** | 2898 |
| Macromolecules | 2796 |
| Ligands | 38 |
| Solvent | 64 |
| **R.m.s. deviation from ideality** | |
| Bond lengths (Å) | 0.010 |
| Bond angles (°) | 1.71 |
| Dihedral angles (°) | 16.8 |
| **Phi-Psi values (Ramachandran)** | |
| Favored (%) | 96.19 |
| Allowed (%) | 3.23 |
| Outliers (%) | 0.59 |
| Rotatmer outliers | 4.61% |
| **Average B-factor** | 54.50 |
| Macromolecules | 54.92 |
| Ligands | 36.01 |
| Solvent | 47.07 |

**Table S2.** SAXS data collection and modeling parameters.

| | SmcR-apo | SmcR-PTSP |
|---|---|---|
| **Data-collection parameters** | | |
| Wavelength (Å) | 1.094 | 1.094 |
| Camera length (m) | 1.755 | 1.755 |
| q-measurement range | 0.0079-0.45 | 0.0079-0.45 |
| Concentration (mg/mL) | 2.01 | 1.24 |
| **Structural parameters** | | |
| I(0) from Guinier | 0.04 ± 0.00145 | 0.03 ± 0.00132 |
| $R_g$ (Å) from Guinier (± SE) | 28.58 ± 0.1 | 30.35 ± 0.13 |
| $R_g$ (Å) from GNOM (± SE) | 29.57 ± 0.57 | 31.58 ± 0.54 |
| I(0) from GNOM | 0.04 ± 0.00345 | 0.03 ± 0.00331 |
| $D_{max}$ (Å) | 125 | 121 |
| $MW^{vp}$ (kDa) | 55.1 | 93.3 |

$R_g$: Radius of gyration. I(0): scattering intensity at zero scattering angle. $D_{max}$: Maximum particle diameter. MW: molecular mass.

**Table S3.** Bacterial strains used in this study.

| Name | Genotype/Notes | Reference |
|---|---|---|
| ***E. coli* strains** | | |
| JV-Ec1008 | SmcR F75Y in pMMB67EH-kanR with pJV064 | This study |
| JV-Ec1011 | SmcR C170F in pMMB67EH-kanR with pJV064 | This study |
| JDN114 | pMMB67EH-kanR empty vector with pJV064 | (1) |
| JDN115 | WT SmcR in pMMB67EH-kanR with pJV064 | (1) |
| TAR-ec001 | SmcR F166C in pMMB67EH-kanR with pJV064 | This study |
| TAR-ec005 | SmcR R179H in pMMB67EH-kanR with pJV064 | This study |
| TAR-ec006 | SmcR F78L in pMMB67EH-kanR with pJV064 | This study |
| TAR-ec007 | SmcR H167Q in pMMB67EH-kanR with pJV064 | This study |
| TAR-ec010 | SmcR R179C in pMMB67EH-kanR with pJV064 | This study |
| TAR-ec012 | SmcR H167R in pMMB67EH-kanR with pJV064 | This study |
| TAR-ec027 | SmcR Q137P in pMMB67EH-kanR with pJV064 | This study |
| TAR-ec040 | SmcR C170Y in pMMB67EH-kanR with pJV064 | This study |
| TAR-ec041 | SmcR I96H in pMMB67EH-kanR with pJV064 | This study |
| TAR-ec042 | SmcR M100L in pMMB67EH-kanR with pJV064 | This study |
| S17-1λpir | Conjugation strain, λ-pir, recA thi pro hsdR- M+ RP4: 2-Tc, Mu, TpR, SmR | (2) |
| DH10B | str. K-12 F– Δ(ara-leu)7697[Δ(rapA'-cra' )] Δ(lac)X74[Δ('yahHmhpE)] duplication (514341-627601) [nmpCgltI] galK16 galE15 e14–(icdWT mcrA) φ80dlacZΔM15 recA1 relA1 endA1 Tn10.10 nupG rpsL150 | Life Technologies |

| | | |
|---|---|---|
| BL21(DE3) | (StrR) rph+ spoT1 Δ(mrr-hsdRMS- mcrBC) λ– Missense (dnaA glmS glyQ lpxK mreC murA) Nonsense (chiA gatZ fhuA? yigA ygcG) Frameshift(flhC mglA fruB)<br><br>E. coli str. B F– ompT gal dcm lon hsdSB(rB–mB–) λ(DE3 [lacI lacUV5-T7p07 ind1 sam7 nin5]) [malB+]K-12(λS) | NEB |
| ***V. vulnificus* strains** | | |
| ATCC 27562 | Wild-type | ATCC |
| CAS-vv015 | ATCC 27562 *ΔsmcR::Spec$^R$*, pMMB67EH-tfoX-kanR | (3) |
| CAS-vv022 | ATCC 27562 SmcR-KmR, pCS32 | (3) |
| CAS-vv026 | ATCC 27562 SmcR-TmR, pCS32 | This study |
| TAR-vv001 | ATCC 27562 SmcR R179C | This study |
| TAR-vv002 | ATCC 27562 SmcR H167Q | This study |
| TAR-vv004 | ATCC 27562 SmcR C170F | This study |
| MO6-24/O | Wild-type | (4) |
| KL321 | MO6-24/O, *ΔclpP* | (4) |
| KL322 | MO6-24/O, *ΔclpA*, KmR | (4) |
| KL421 | MO6-24/O, *Δlon*, KmR | (4) |
| SM801 | MO6-24/O, *ΔclpX*, KmR | (4) |
| TAR-vv101 | MO6-24/O, pCS38 | This study |
| TAR-vv103 | MO6-24/O, *ΔclpA,* pCS38 | This study |
| TAR-vv105 | MO6-24/O, *ΔclpX,* pCS38 | This study |
| TAR-vv107 | MO6-24/O, *ΔclpP,* pCS38 | This study |
| TAR-vv109 | MO6-24/O, *Δlon,* pCS38 | This study |
| TAR-vv117 | ATCC 27562, *ΔclpP* | This study |
| TAR-vv119 | ATCC 27562, *ΔclpA* | This study |
| TAR-vv121 | ATCC 27562, *ΔclpX* | This study |
| TAR-vv111 | ATCC 27562 SmcR R179C*,* pCS38 | This study |
| TAR-vv113 | ATCC 27562 SmcR H167Q*,* pCS38 | This study |
| TAR-vv115 | ATCC 27562 SmcR C170F*,* pCS38 | This study |
| ADP-vv001 | ATCC 27562, *ΔclpP,* pCS38 | This study |
| ADP-vv002 | ATCC 27562, *ΔclpA,* pCS38 | This study |
| ADP-vv003 | ATCC 27562, *ΔclpX,* pCS38 | This study |
| FA-021 | ATCC 27562, *ΔluxO::TmR*, pCS38 | This study |
| FA-055 | ATCC 27562, *luxO D60E,* pCS38 | This study |

| | | |
|---|---|---|
| ***V. campbellii* strains**<br>BB120 | Wild-type | (5) |
| KM669 | BB120 *ΔluxR* | (6) |

**Table S4.** Oligonucleotides used in this study.

| Name | Direction | Sequence | Description |
|---|---|---|---|
| TAR001 | F | GACAATTAATCATCGGCTCGTATAATG | pJN22 sequencing, F |
| TAR002 | R | GCGTTCTGATTTAATCTGTATCAGGC | pJN22 sequencing, R |
| TAR004 | F | CAGAGAAACGCTAACTCAGGTACGCGGATG GAGGAGGGCCCTAGGAC | Primer to amplify SmcR insert from WT ATCC 27562 |
| TAR007 | R | CTCATCCGCCAAAACAGCCAAGCTCTATTC GTGTTCGCGTTTATAGATG | Primer to amplify SmcR insert from WT ATCC 27562 |
| TAR010 | F2 | TCGCTGTTTGTTCAAGCAAACTGCACCAACA ATACTGCAGAGCTA | F2 primer SmcR R179C |
| TAR011 | R1 | TAGCTCTGCAGTATTGTTGGTGCAGTTTGCT TGAACAAACAGCGA | R1 primer SmcR R179C |
| TAR012 | R | CTCAGGTCGACGGATCCCCGGAAT | R sequencing primer to check SmcR mutations in CAS-vv022 |
| TAR015 | F2 | GAAGATTTGGCAAACCTGTTCCAAGGCATTT GTTACTCGCTGTTT | F2 primer SmcR H167Q |
| TAR016 | R1 | AAACAGCGAGTAACAAATGCCTTGGAACAG GTTTGCCAAATCTTC | R1 primer SmcR H167Q |
| JCV1311 | F2 | ATTTGGCAAACCTGTTCCACGGCATTTTCTA CTCGCTGTTTGTTCAAGCAAACCGC | F2 primer SmcR C170F |
| JCV1312 | R1 | GCGGTTTGCTTGAACAAACAGCGAGTAGAA AATGCCGTGGAACAGGTTTGCCAAAT | R1 primer SmcR C170F |
| CAS107 | F1 | CGTCTCAGGCAACGCAGACAGTG | pMMB amplify backbone, F |
| CAS110 | R2 | GGAAGCTCAAGCAACGACTAGTG | pMMB amplify backbone, R |
| TAR029 | F1 | AATCGTCTCAGGCAACGCAGACAGTGTAC | Redesigned CAS107 |
| TAR030 | R2 | AGCGGAAGCTCAAGCAACGACTAGTGATT | Redesigned CAS110 |
| TAR044 | F1 | AGCGTCGAACTCCACCCCAGTTGG | F1, clpP deletion in WT ATCC 27562 |
| TAR045 | R1 | GTCGACGGATCCCCGGAATACTCATCGTAG TGCATAATTTGCCTGC | R1 with homology to ABD123, clpP deletion in WT ATCC 27562 |
| TAR046 | F2 | GAAGCAGCTCCAGCCTACATAAGCTGGTTG GTTCATTAGCTCG | F2 with homology to ABD124, clpP deletion in WT ATCC 27562 |
| TAR047 | R2 | GAACTTCGCAGGATCGATGCCTGG | R2, clpP deletion in WT ATCC 27562 |
| TAR048 | R | GAACTAAACGACGAGTTCGTT | ΔclpP detection primer reverse |
| TAR049 | F1 | CGCTAGCAACTTACTCATTGCTGCC | F1, clpA deletion in WT ATCC 27562 |
| TAR050 | R1 | GTCGACGGATCCCCGGAATTATTAAGCATA AGTACCTCCTGTG | R1 with homology to ABD123, clpA deletion in WT ATCC 27562 |

| | | | |
|---|---|---|---|
| TAR051 | F2 | GAAGCAGCTCCAGCCTACAAGCGGTTCATT AAAAGTAAATAAT | F2 with homology to ABD124, clpA deletion in WT ATCC 27562 |
| TAR052 | R2 | CAACCCTAACCCCATCATCAA | R2, clpA deletion in WT ATCC 27562 |
| TAR053 | R | GAATACGCCCCAACTTTAC | ΔclpA detection primer reverse |
| TAR054 | F | CCGGCACTTGGGCCATCTTTT | F1, clpX deletion in WT ATCC 27562 |
| TAR055 | R | GTCGACGGATCCCCGGAATCGCTGGCGCT GAGTAAATTTAATC | R1 with homology to ABD123, clpX deletion in WT ATCC 27562 |
| TAR056 | F | GAAGCAGCTCCAGCCTACATGCTTTTGTCT GTCATTCGCTAAC | F2 with homology to ABD124, clpX deletion in WT ATCC 27562 |
| TAR057 | R | GTTTCATGGCGAAACCGATAC | R2, clpX deletion in WT ATCC 27562 |
| TAR058 | R | CGAAGATGGTATCGAAGG | ΔclpX detection primer reverse |
| ABD123 | R | ATTCCGGGGATCCGTCGAC | Forward primer to amplify antibiotic resistance cassettes |
| ABD124 | F | TGTAGGCTGGAGCTGCTTC | Reverse primer to amplify antibiotic resistance cassettes |
| JCV 149 | F | CGTGTCGCTCAAGGCGCACTCCCGT | pJN22 amplify backbone, F |
| JDN104 | R | ATCGAGTTGCGGCCGCTTGTTGTTACCTTA GCAGGGTG | pJN22 amplify backbone, R |
| CAS536 | R | CAACCAACAGCCATTGGTTATAGG | R to detect in TmR cassette for seq before TmR |

**Table S5.** Plasmids used in this study.

| **Name** | **Description** | **Reference** |
|---|---|---|
| pJN08 | 6x-his-smcR in pET28b | (1) |
| pJN22 | $P_{tac}$-smcR (V. vulnificus ATCC 27562), in pMMB67EH-kanR | (1) |
| pJV064 | $P_{05222}$-mCherry, $P_{luxC}$-gfp; p15a origin, $CM^R$ | (8) |
| pCS38 | *luxCDABE* region, in pMMB67EH-kanR | (1) |
| pMMB67EH-kanR | Vector control; kanR | (3) |
| pTR001 | *His-smcR C170F*; pET28a parent vector | This study, TWIST |
| pTR002 | *His-smcR F75Y*; pET28a parent vector | This study, TWIST |
| pTR003 | *His-smcR R179C*, pET28a.parent vector | This study, TWIST |

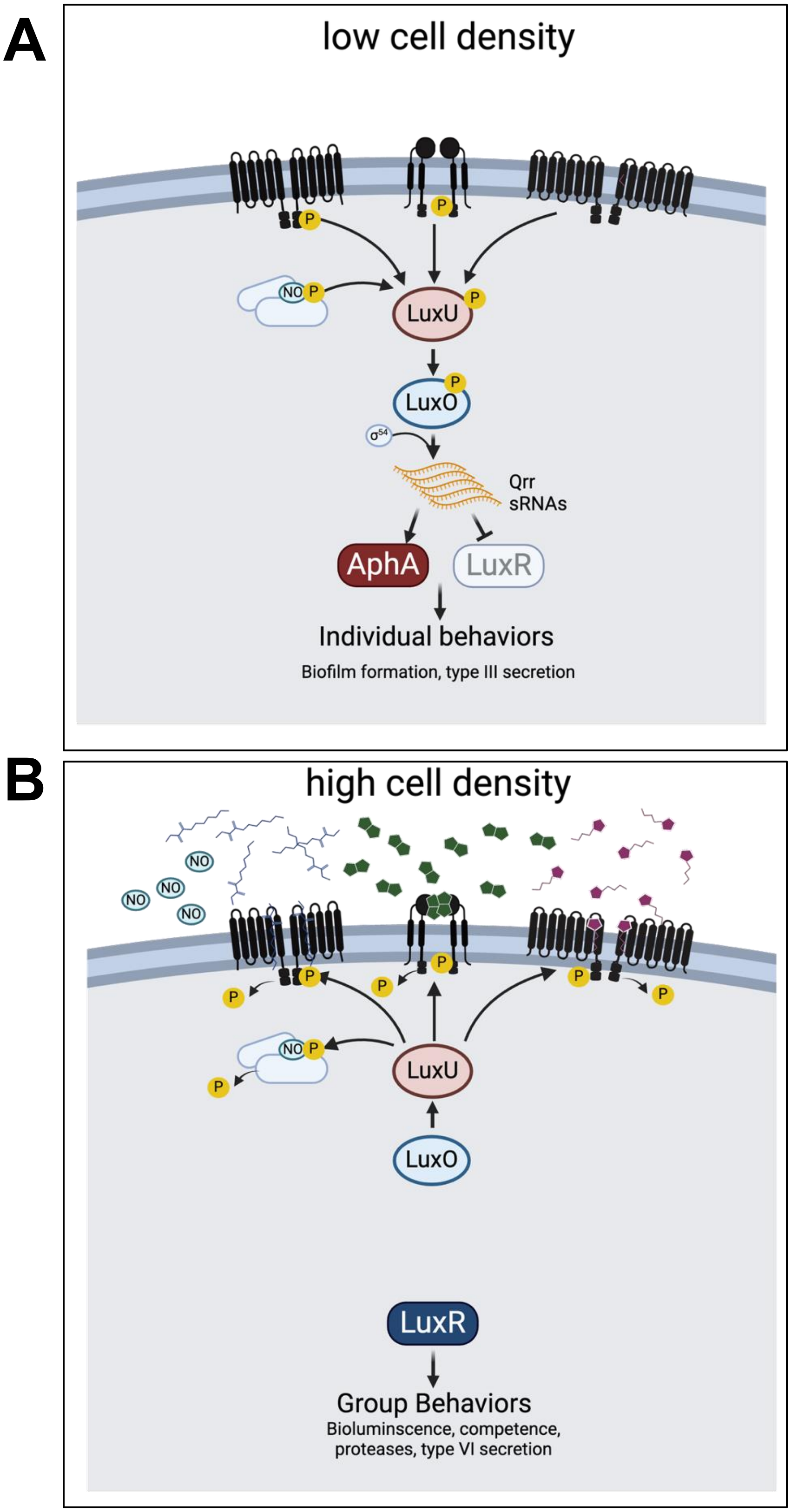

**Figure S1. Current model of QS signaling at low and high cell density in *V. campbellii*.** (A) At low cell density, autoinducer concentrations are low and do not bind to receptors. Receptors act as kinases to phosphorylate (P) LuxU and subsequently LuxO. Phosphorylated LuxO and Sigma-54 activate transcription of the Qrr sRNAs, which activate translation of AphA and repress translation of LuxR, leading to transcriptional regulation of genes driving individual behaviors. (B) At high cell density, high levels of autoinducers and/or nitric oxide (NO) enable binding to the receptors, which results in phosphatase activity. Dephosphorylated LuxO does not activate the Qrrs, and thus LuxR is translated to high levels. Transcription by LuxR regulates hundreds of genes required for producing group behaviors.

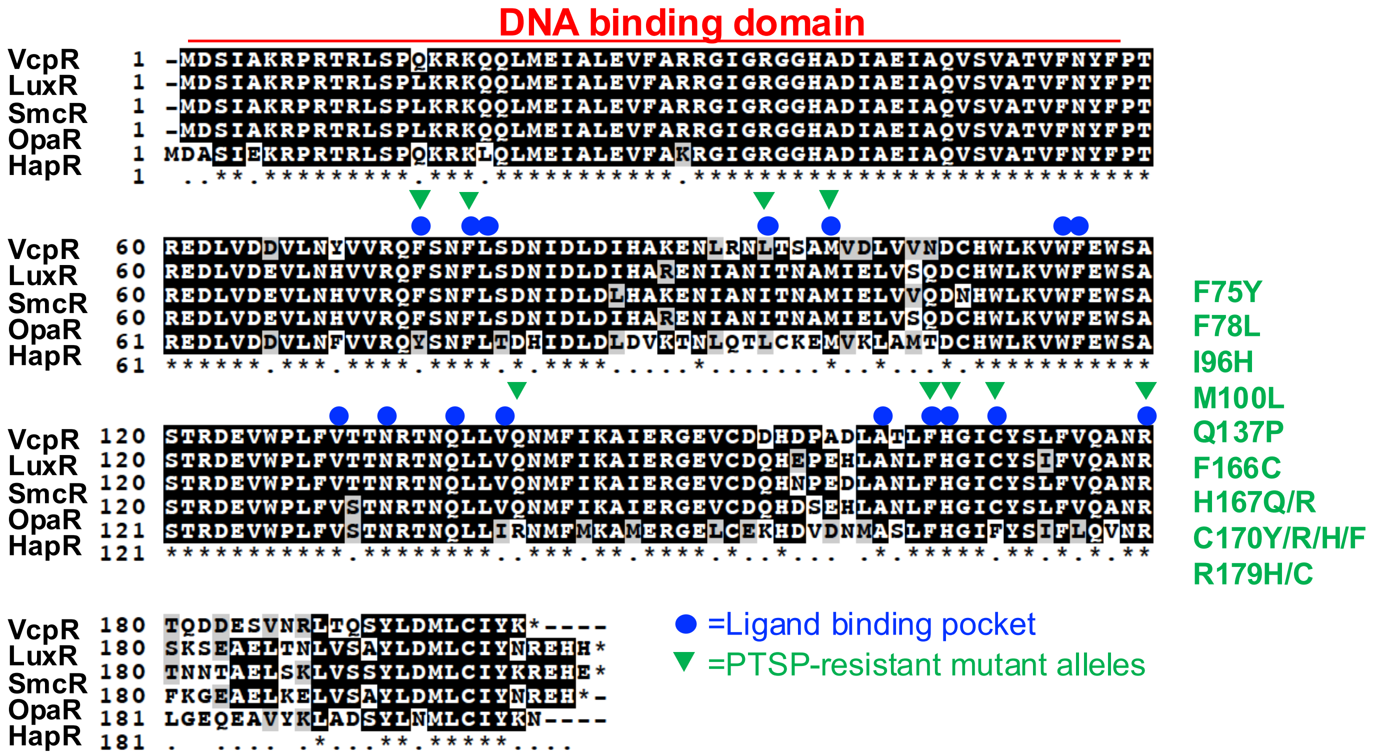

**Figure S2. Alignment of the LuxR/HapR homologues.** VcpR (*Vibrio coralliilyticus* OCN008), LuxR (*V. campbellii* BB120), SmcR (*V. vulnificus* ATCC 27562), OpaR (*V. parahaemolyticus* RIMD2210633), and HapR (*V. cholerae* E7946). The residues identified to be part of the ligand binding pocket of SmcR-PTSP are indicated by a blue dot above the alignment position. The residues identified as PTSP-resistant alleles by genetics are indicated by green triangles above the alignment position.

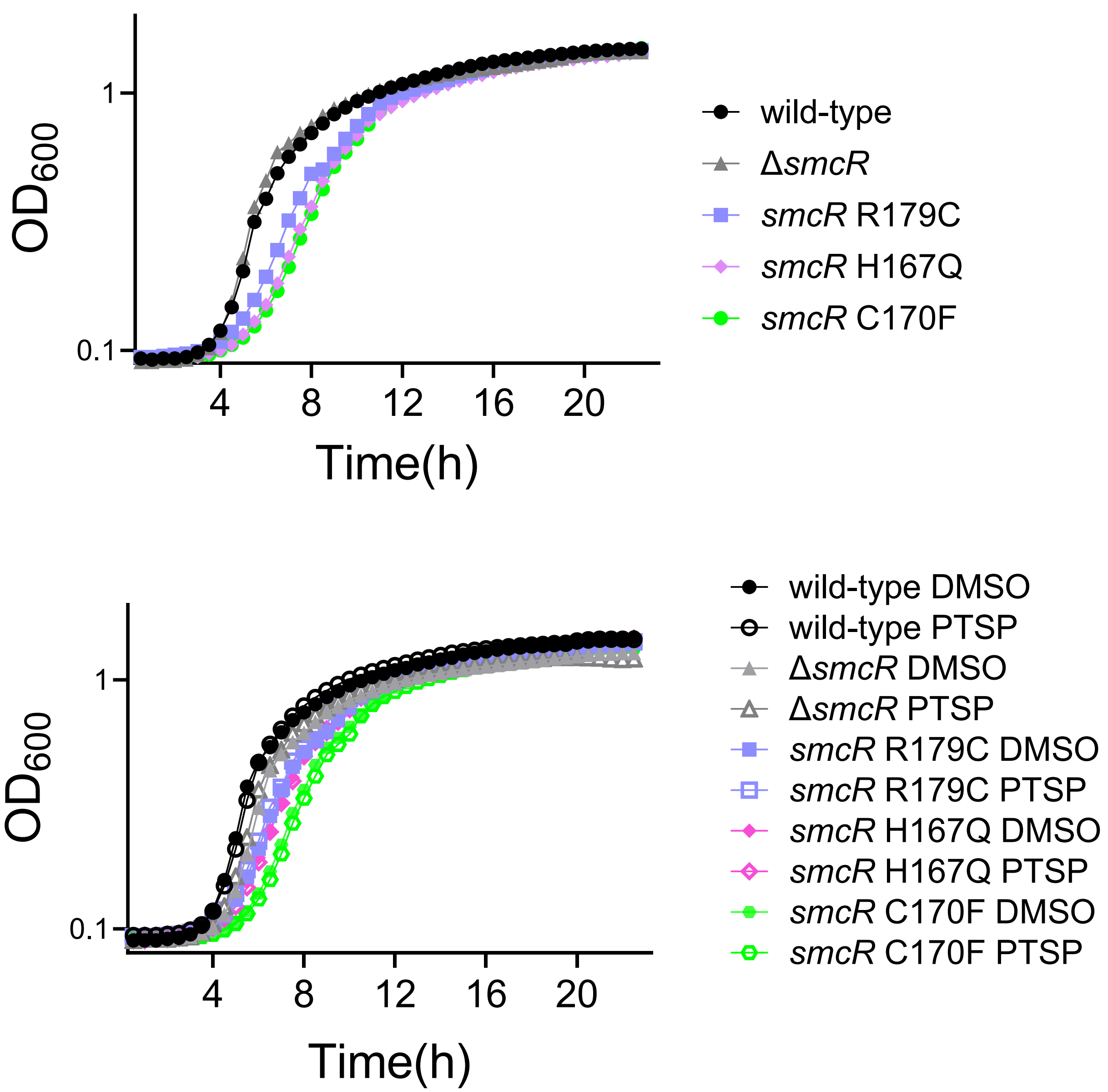

**Figure S3. Addition of PTSP to *Vibrio vulnificus* does not alter growth trends.** (A) Overnight growth curve assays of SmcR protein in V*ibrio vulnificus* strains expressing wild-type *smcR,* Δ*smcR*, *smcR* R179C, H167Q, or C170F treated with 25 μM PTSP or an equivalent concentration of DMSO solvent. Figures are representative of triplicate biological assays

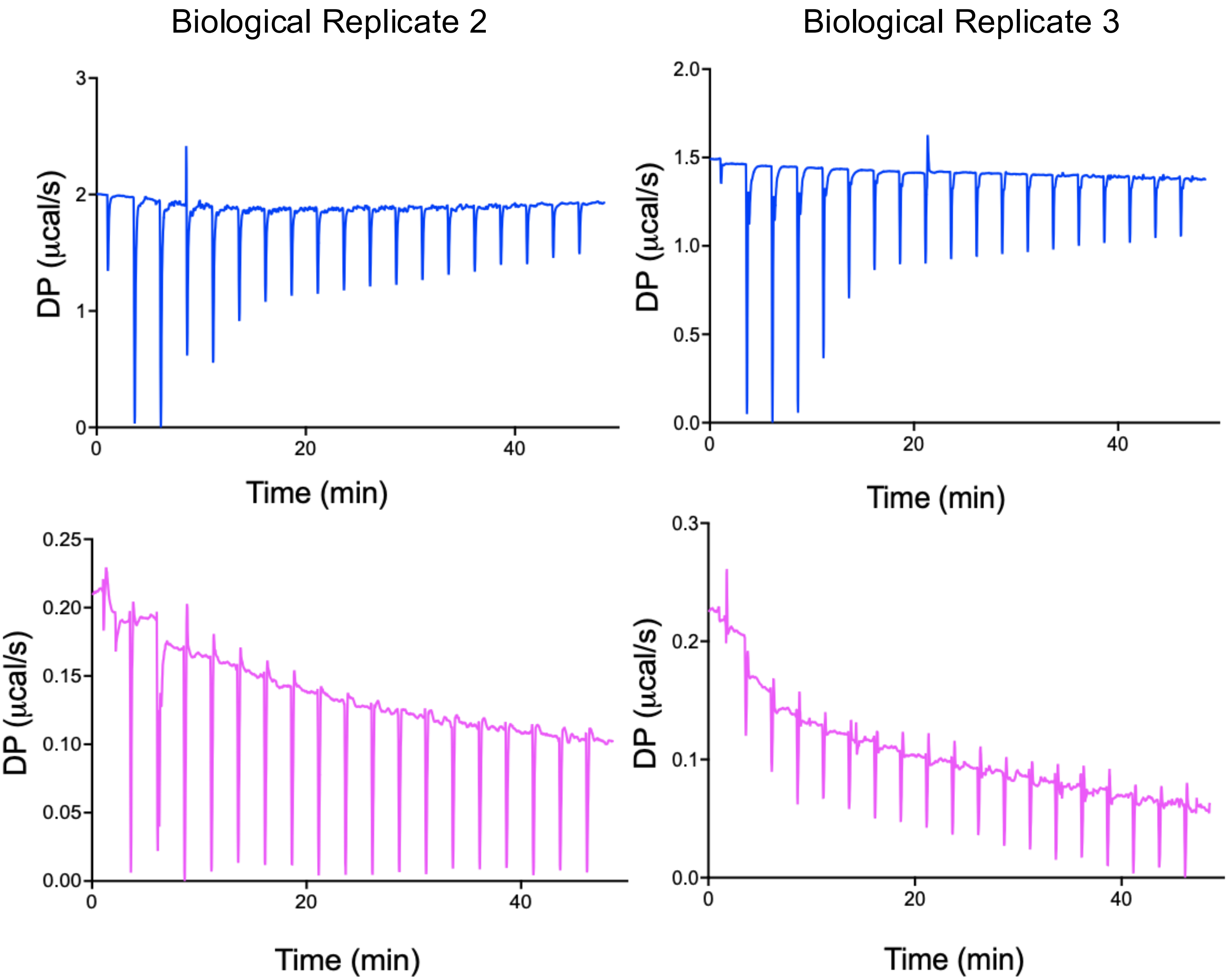

**Figure S4. Isothermal titration calorimetry of wild-type SmcR and SmcR R179C.** Biological replicates of isothermal titration calorimetry experiments of WT SmcR (blue) and SmcR R179C (magenta) with PTSP for Figure 1. DP denotes differential power. Different protein purification preparations were used for each replicate.

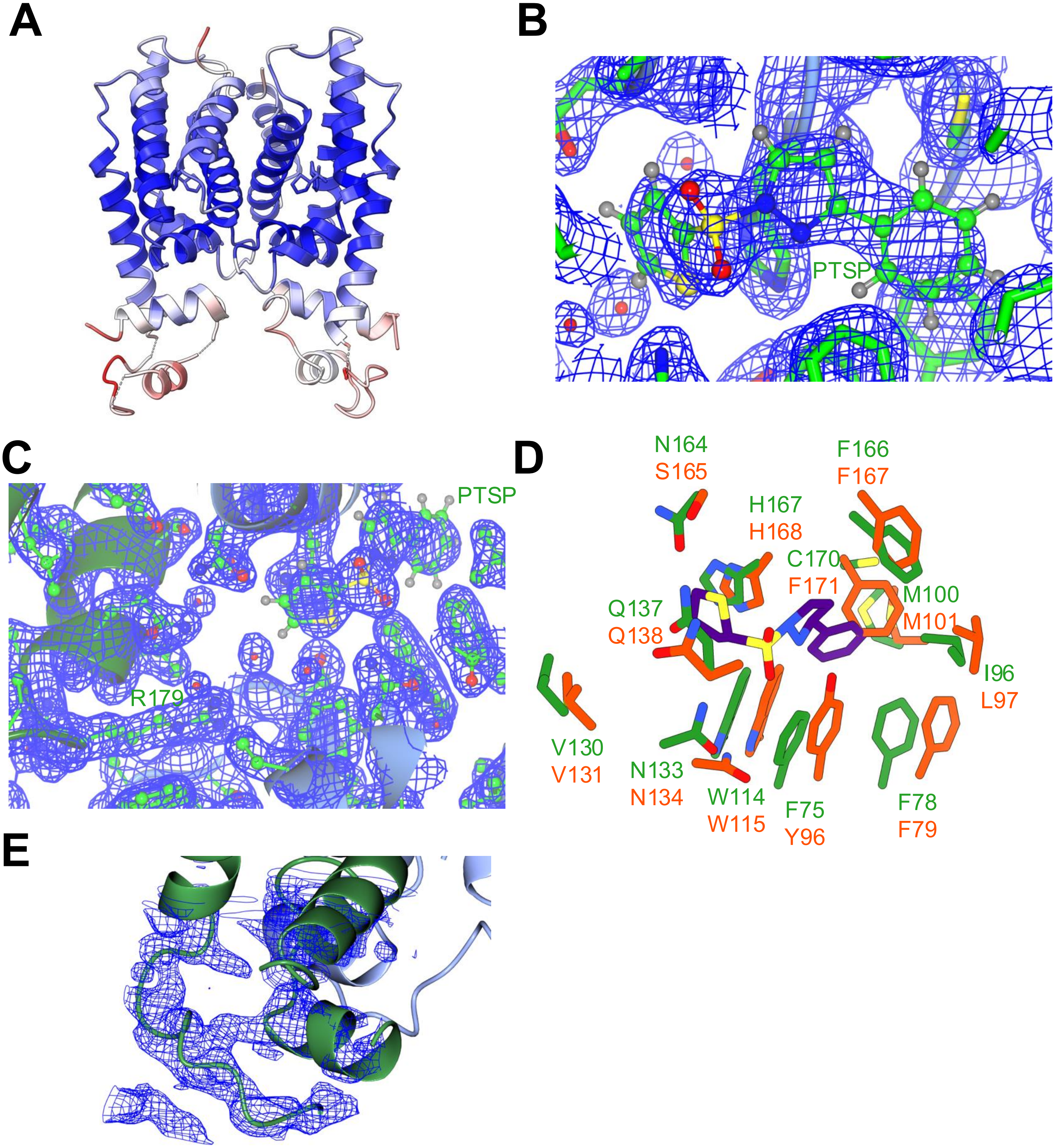

**Figure S5. Validation of the SmcR-PTSP structure.** (A) Structure of SmcR-PTSP colored by B-factor. Blue indicates low B-factor and red indicates high B-factor scores. (B) Example close-up view of map density of the SmcR LBP-SmcR interface and (C) R179. (D) Structural overlay of the residues important for ligand binding in SmcR (green) and HapR (orange) in their respective binding pocket relative to the orientation of PTSP (purple) in SmcR. (E) Example close-up view of map density of the SmcR N-terminal domain.

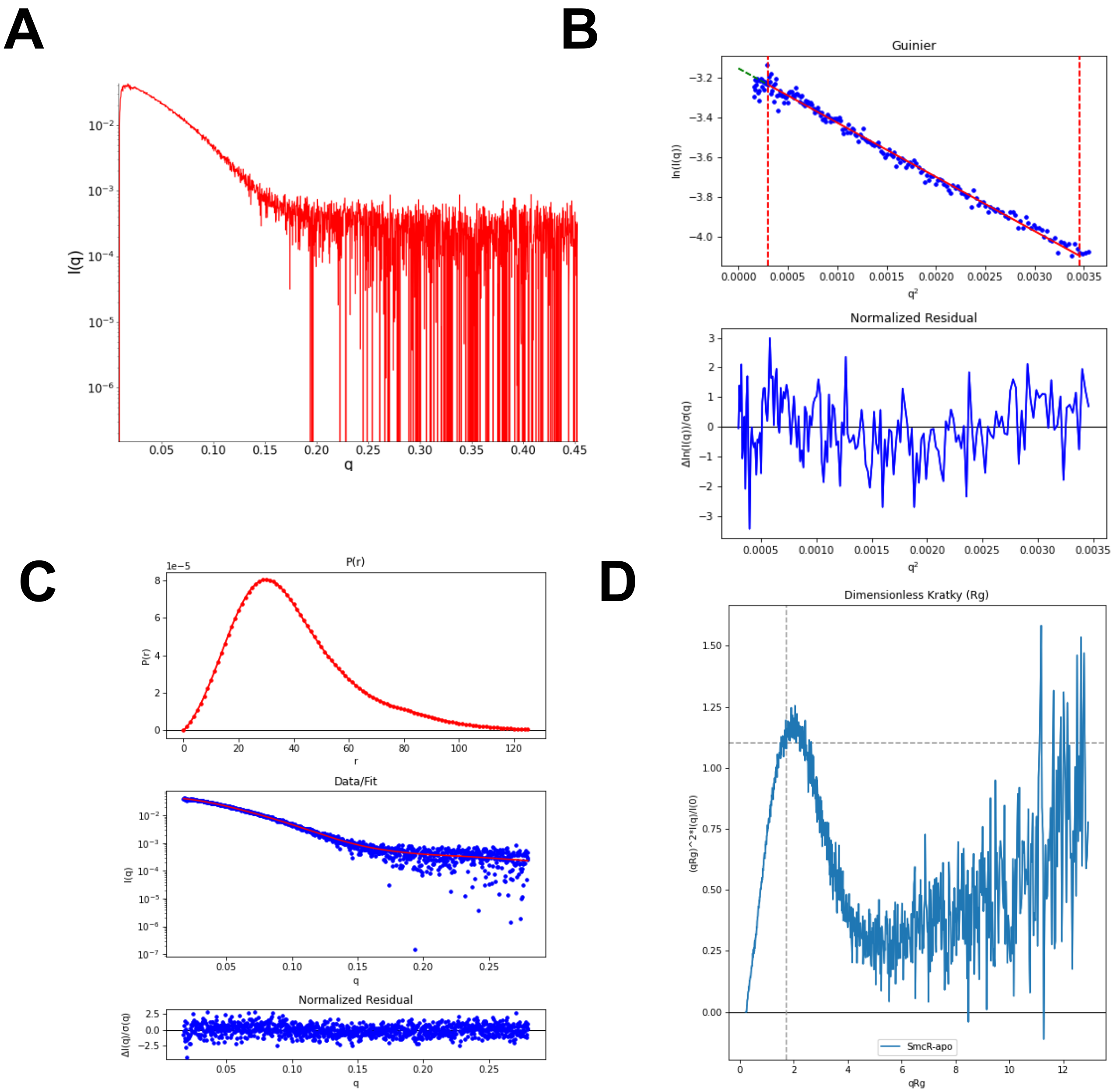

**Figure S6. SAXS summary plots for SmcR-apo.** (A) Scattering intensity profile, I(q) as a function of the scattering vector (q) on a log-lin scale. (B) Guinier plot (top panel) and normalized residuals (bottom panel). (C) The pair-distance distribution function P(r) (top panel). The scattering intensity profile calculated from the P(r) function (red) overlaid with the scattering intensity profile from the experimental data (blue, middle panel). (D) Dimensionless Kratky plot. Dashed lines show where a globular system would peak.

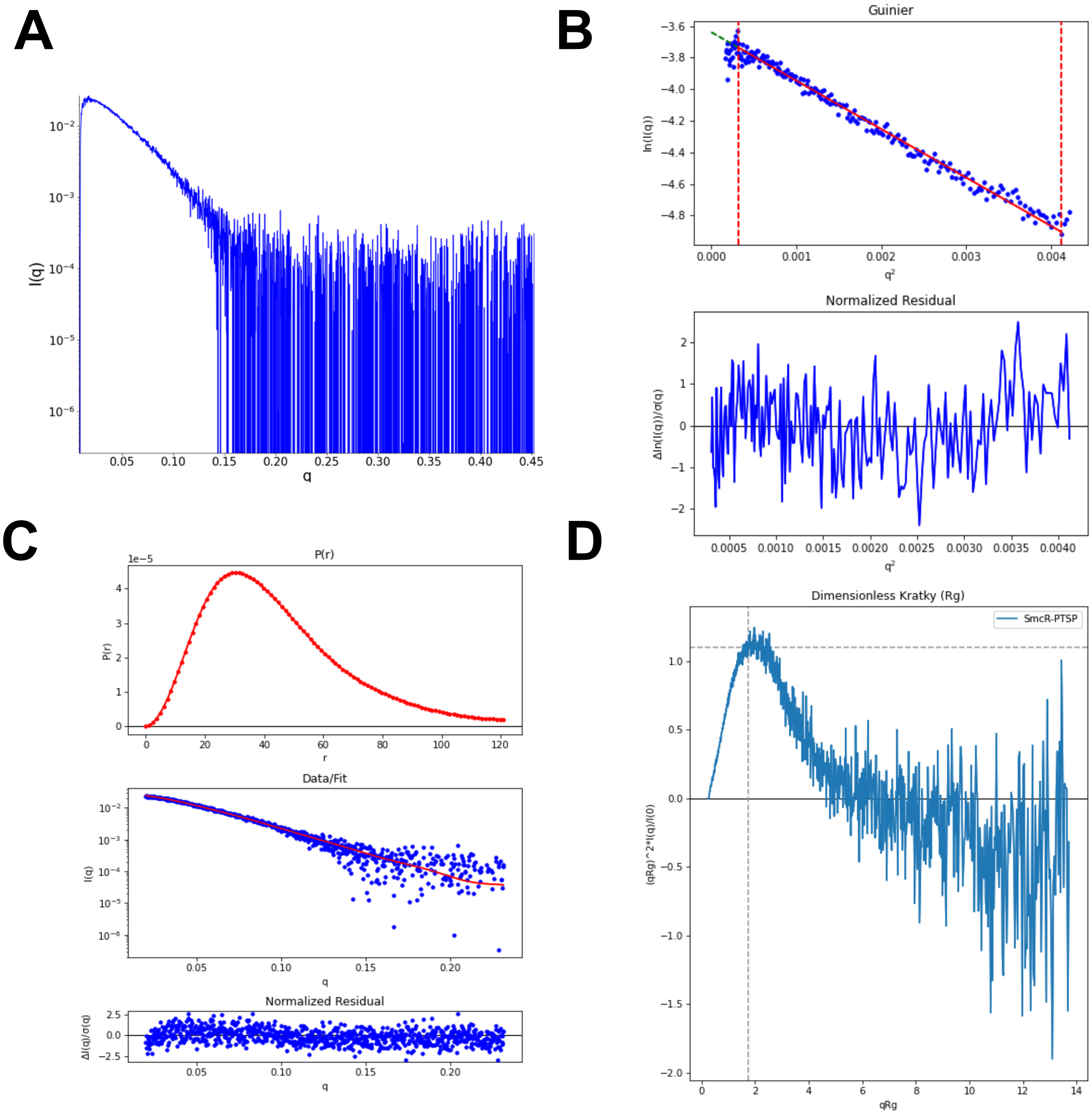

**Figure S7. SAXS summary plots for SmcR-PTSP.** (A) Scattering intensity profile, I(q) as a function of the scattering vector (q) on a log-lin scale. (B) Guinier plot (top panel) and normalized residuals (bottom panel). (C) The pair-distance distribution function P(r) (top panel). The scattering intensity profile calculated from the P(r) function (red) overlaid with the scattering intensity profile from the experimental data (blue, middle panel). (D) Dimensionless Kratky plot. Dashed lines show where a globular system would peak.

A

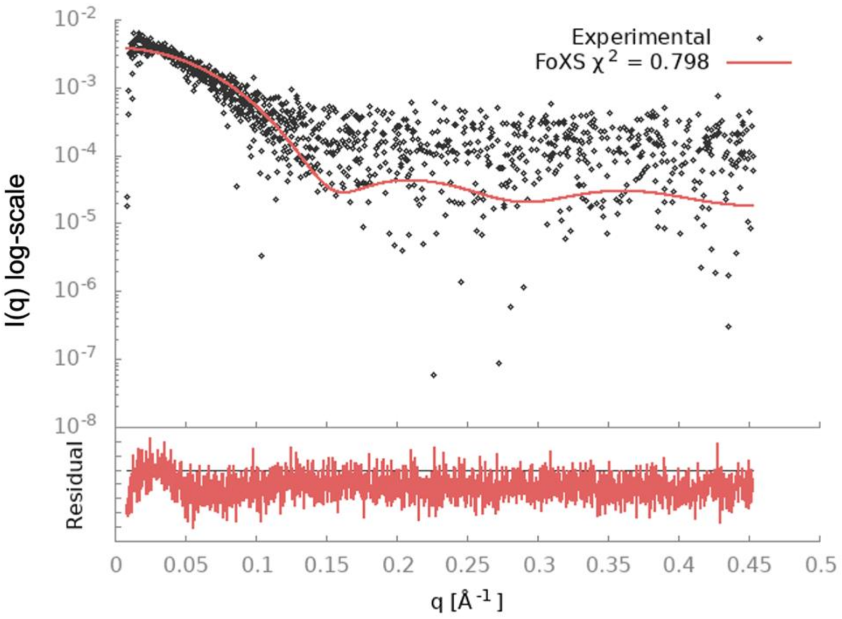

B

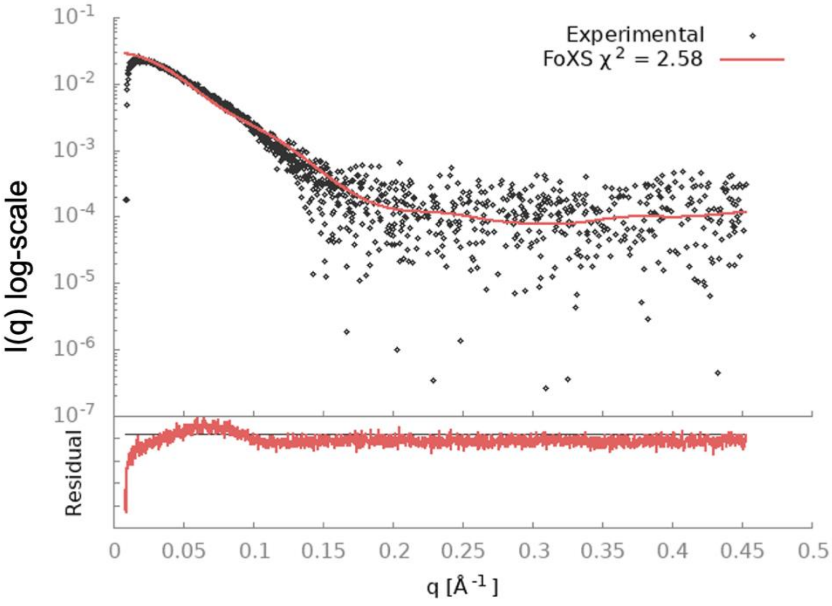

**Figure S8. Single-state modeling of SmcR proteins based on experimental SAXS profiles*.*** (A) The SmcR-apo SAXS experimental data (black circles) was fit to the theoretical profile derived from the high-resolution X-ray structure (PDB: 6WAE) (red line). (B) The SmcR-PTSP SAXS experimental data (black circles) was fit to the theoretical profile derived from the high-resolution X-ray structure (PDB: 6WAE) (red line). Single-state modeling was performed using the FoXS server (https://modbase.compbio.ucsf.edu/foxs/) (Schneidman-Duhovny et al., 2013; Schneidman-Duhovny et al., 2016).

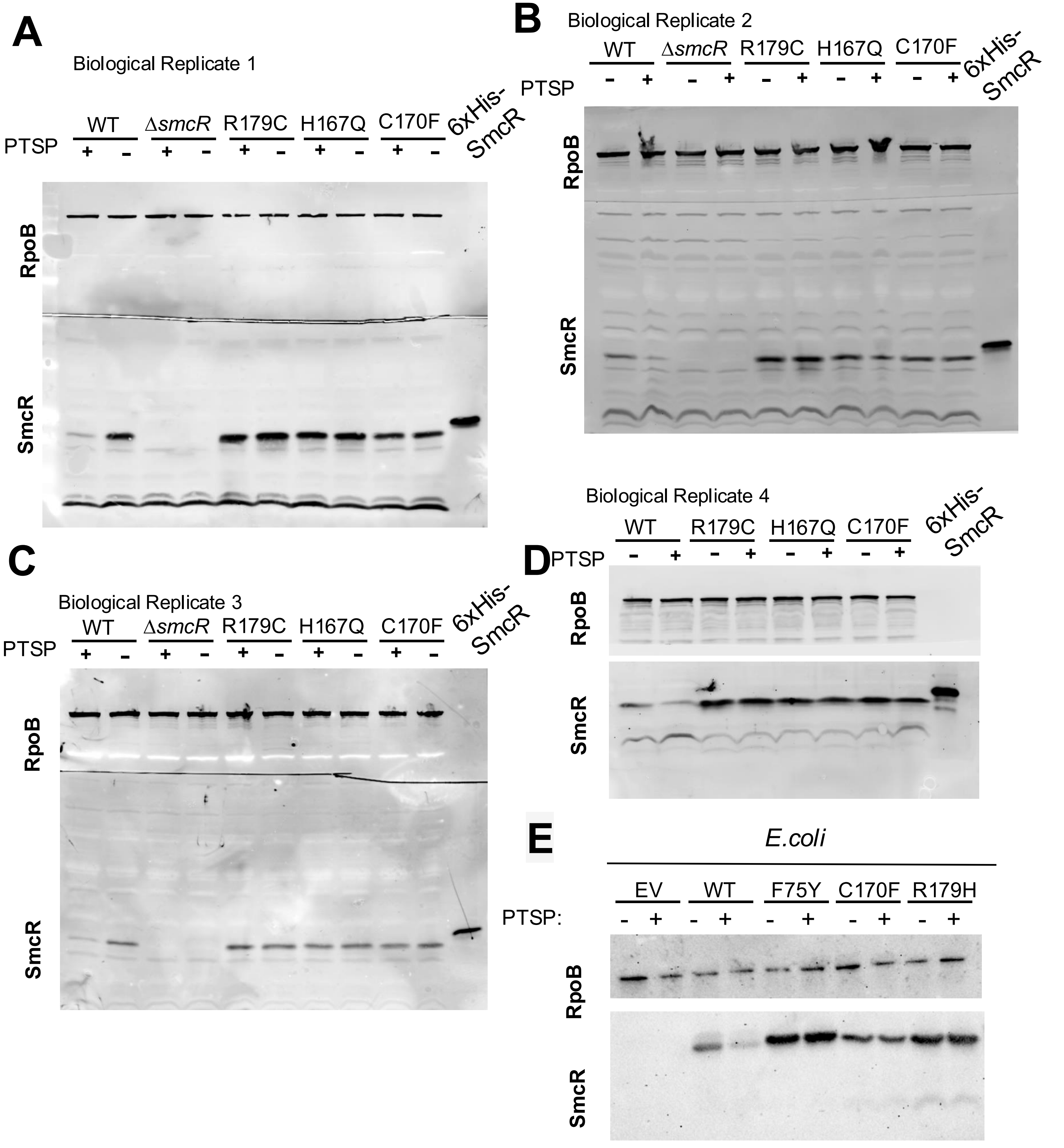

**Figure S9. PTSP increases proteolytic degradation of SmcR *in vivo.*** (A-D) Western blot analysis of SmcR protein levels in cell lysates from *V. vulnificus* strains wild-type, *smcR* R179C, H167Q and C170F with 25 μM PTSP (+) or an equivalent volume of DMSO solvent (-). Anti-RpoB antibodies were used as a loading control. Each panel is one biological replicate. Uncropped gels are shown. (E) Western blot analysis of SmcR protein levels in cell lysates from *E.coli* strains expressing wild-type *smcR*, *smcR* F75Y, C170F, R179H, or containing an empty vector (EV) treated with 25 μM PTSP (+) or an equivalent concentration of DMSO solvent (-). Anti-RpoB antibodies were used as a loading control.

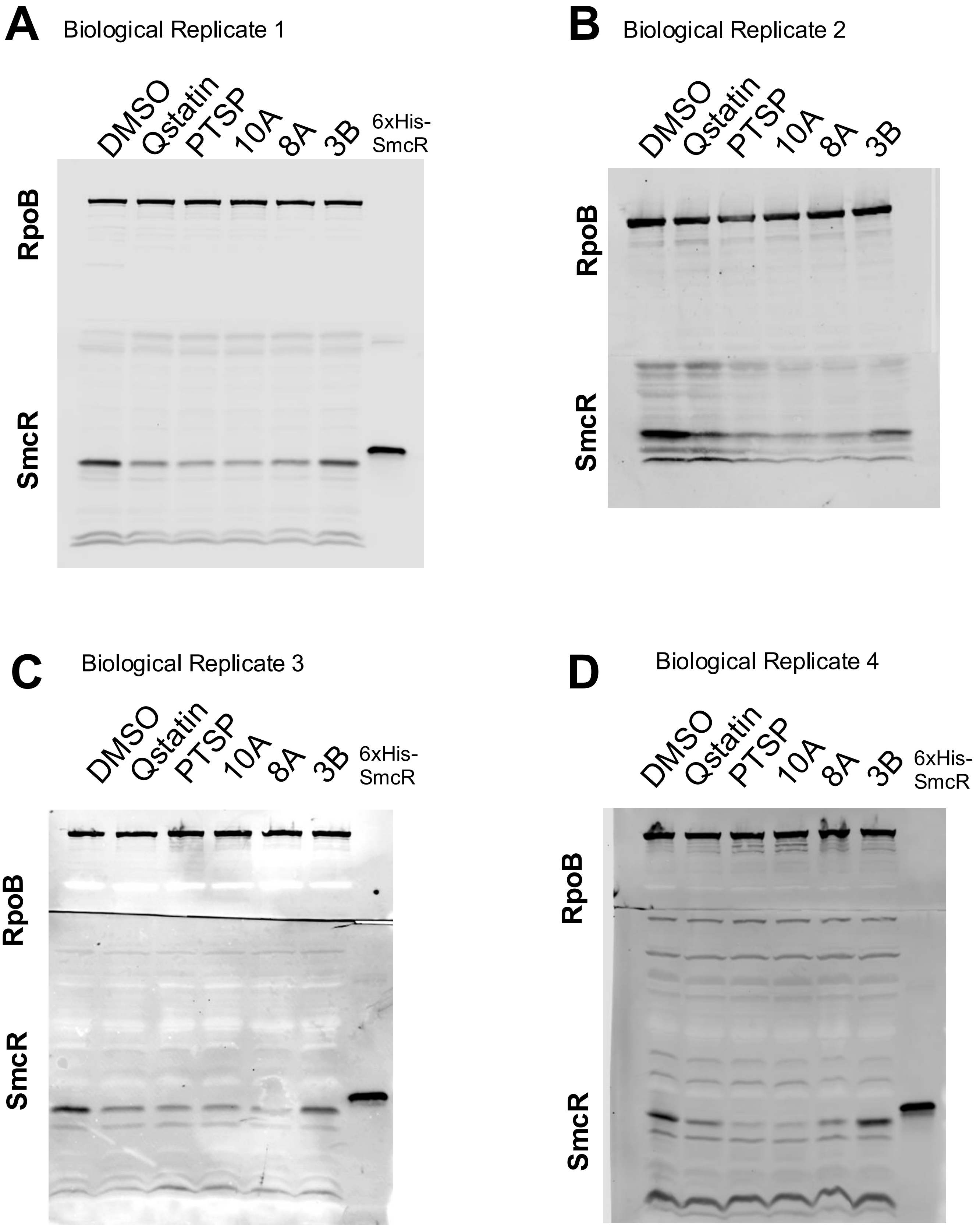

**Figure S10.** (A-C) Western blot analysis of SmcR protein levels in cell lysates from wild-type *V. vulnificus* treated with 25 $\mu$M of each thiophenesulfonamide or an equivalent volume of DMSO solvent. Anti-RpoB antibodies were used as a loading control. Each panel is one biological replicate. Uncropped gels are shown.

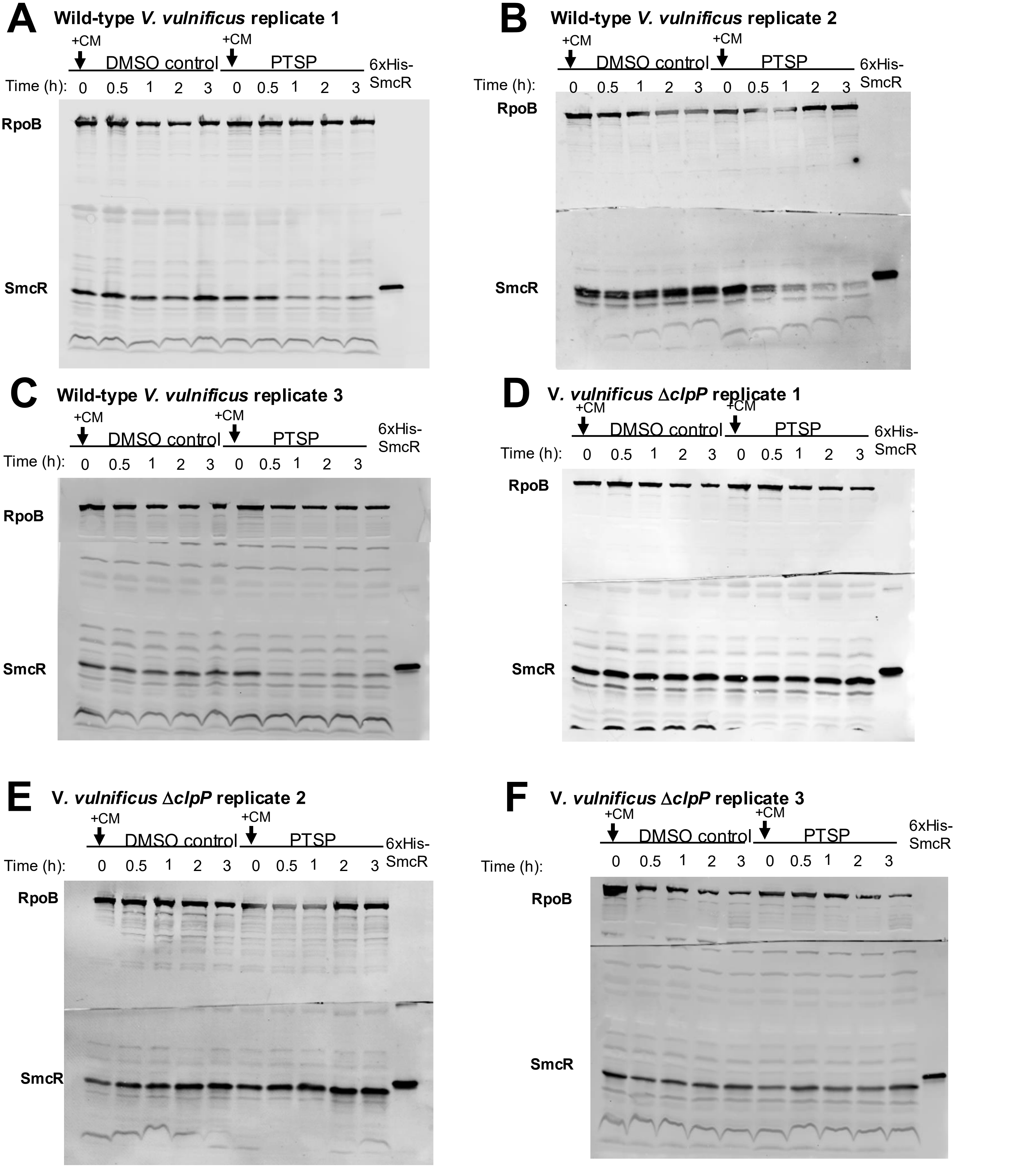

**Figure S11. Time-course of SmcR protein stability following translation inhibition.** (A-F) Western blot analysis of cell lysates with anti-LuxR antibodies measuring SmcR from *V. vulnificus,* wild-type (A-C) or Δ*clpP* (D-F). Cells were either treated with 25 μM PTSP or an equivalent concentration of DMSO solvent and grown to mid-log phase ($OD_{600}$ = 0.5-0.8). Translation was halted by addition of 10 μg/ml of chloramphenicol (CM) at time 0, and pellets were harvested at 0, 30, 60, 120, and 180 mins. Anti-RpoB antibodies were used as a loading control. Each panel is one biological replicate. Uncropped gels are shown.

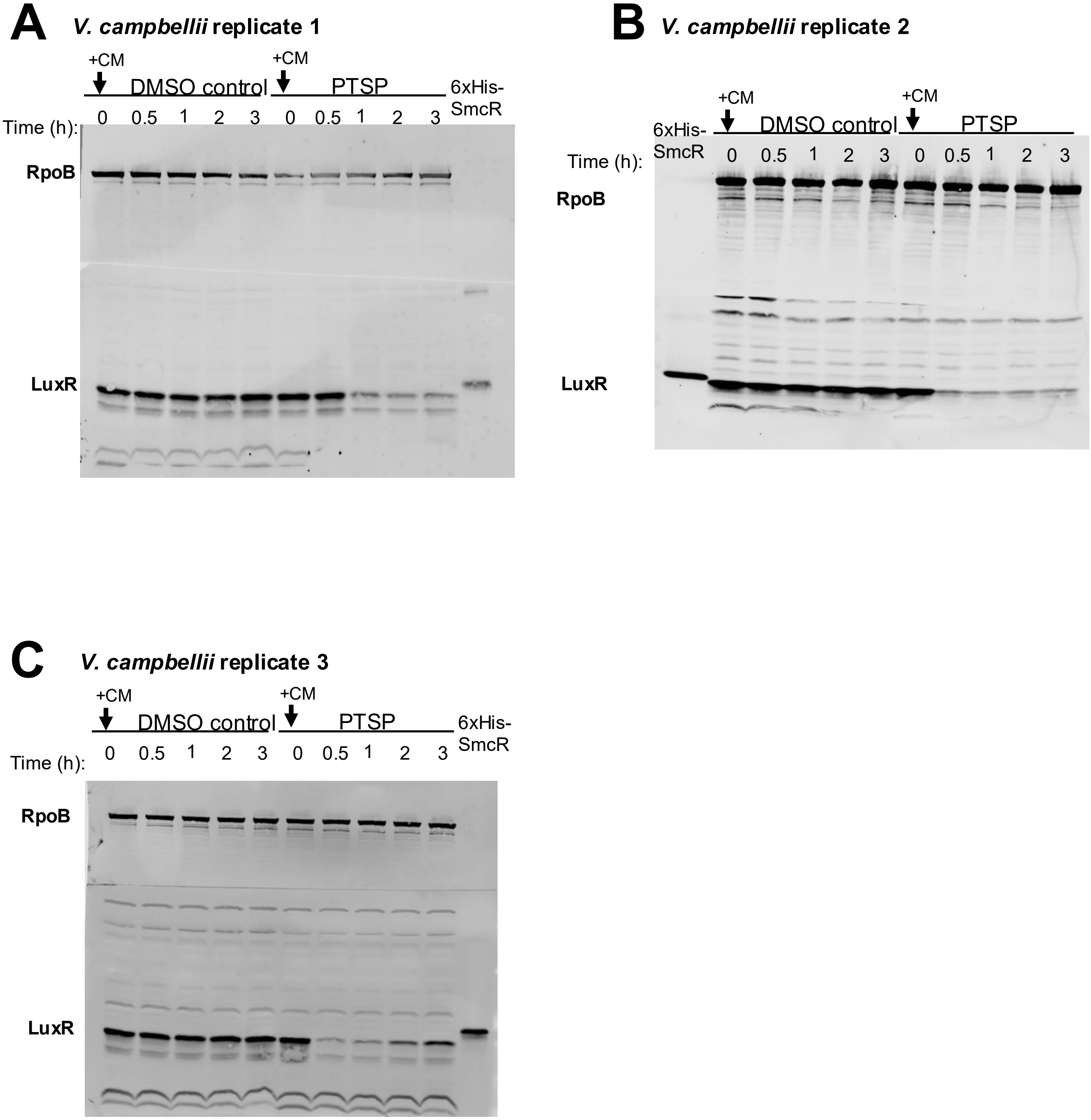

**Figure S12. Time-course of LuxR protein stability following translation inhibition.** (A, B, C) Western blot analysis of cell lysates with anti-LuxR antibodies measuring LuxR from *V. campbellii.* Cells were either treated with 25 μM PTSP or an equivalent concentration of DMSO solvent and grown to mid-log phase ($OD_{600}$ = 0.5-0.8). Translation was halted by addition of 10 μg/ml of chloramphenicol (CM) at time 0, and pellets were harvested at 0, 30, 60, 120, and 180 mins. Anti-RpoB antibodies were used as a loading control. Each panel is one biological replicate. Uncropped gels are shown.

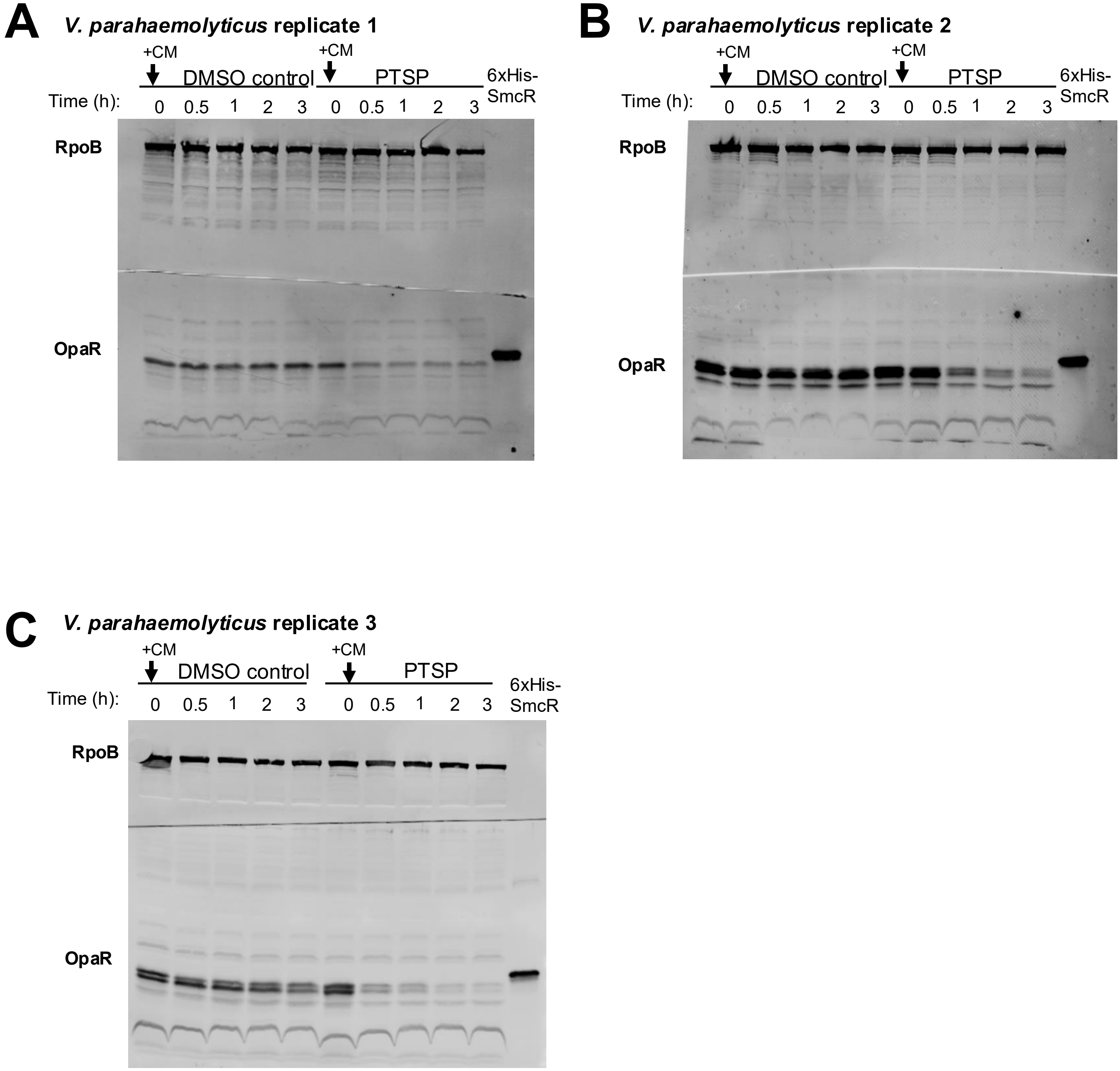

**Figure S13. Time-course of OpaR protein stability following translation inhibition.** (A, B, C) Western blot analysis of cell lysates with anti-LuxR antibodies measuring OpaR from *V. parahaemolyticus.* Cells were either treated with 25 μM PTSP or an equivalent concentration of DMSO solvent and grown to mid-log phase ($OD_{600}$ = 0.5-0.8). Translation was halted by addition of 10 μg/ml of chloramphenicol (CM) at time 0, and pellets were harvested at 0, 30, 60, 120, and 180 mins. Anti-RpoB antibodies were used as a loading control. Each panel is one biological replicate. Uncropped gels are shown.

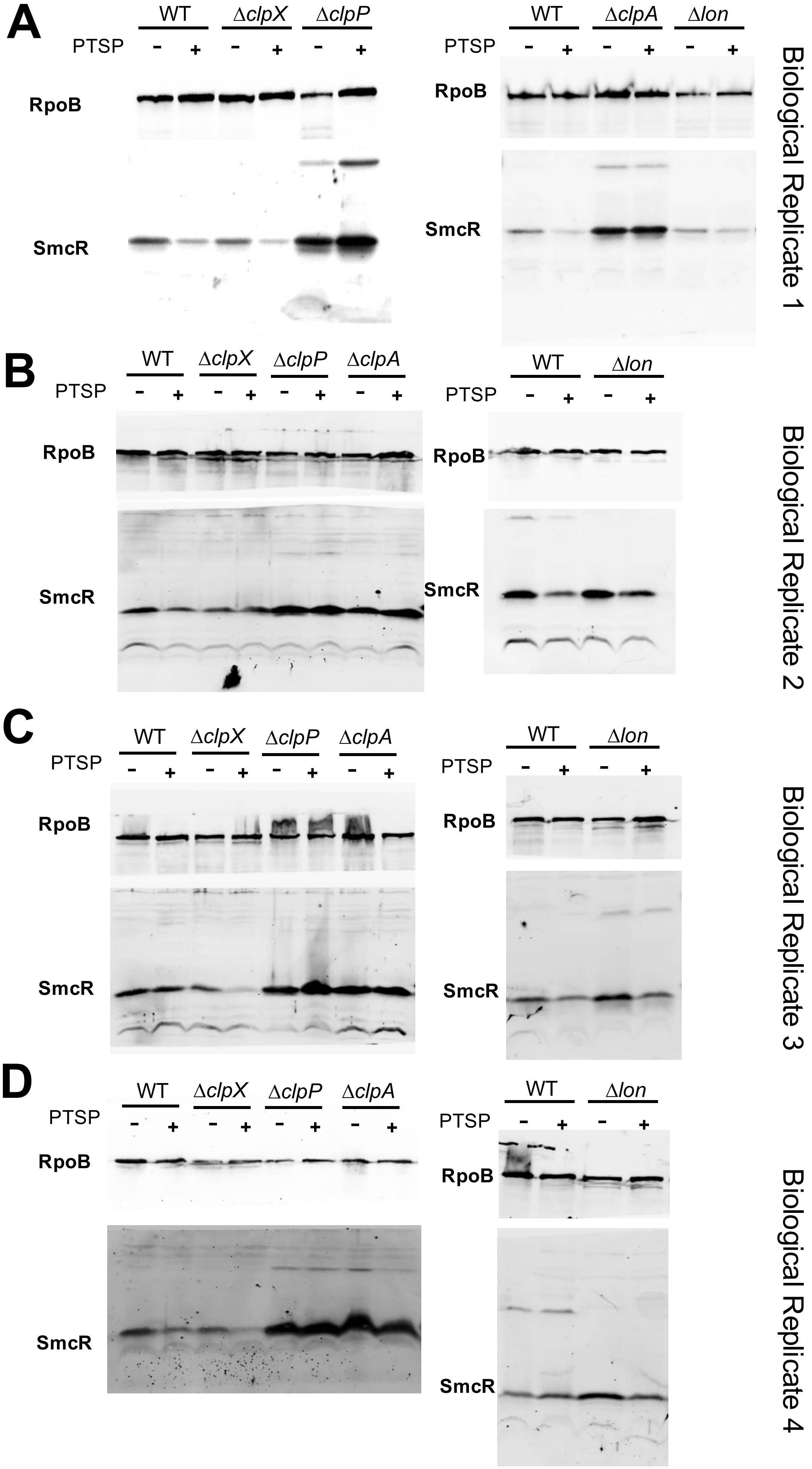

**Figure S14. ClpAP proteolysis of SmcR is blocked by substitutions that block inhibitor binding.** Western blot analysis of cell lysates measuring SmcR from *V. vulnificus* strain MO6-24/0 wild-type, Δ*clpX*, Δ*clpP*, or Δ*clpA*, or Δ*lon*. Cells were treated with 25 μM of PTSP or an equivalent concentration of DMSO solvent (-). Anti-RpoB antibodies were used as a loading control. Each panel is one biological replicate. Uncropped gels are shown.

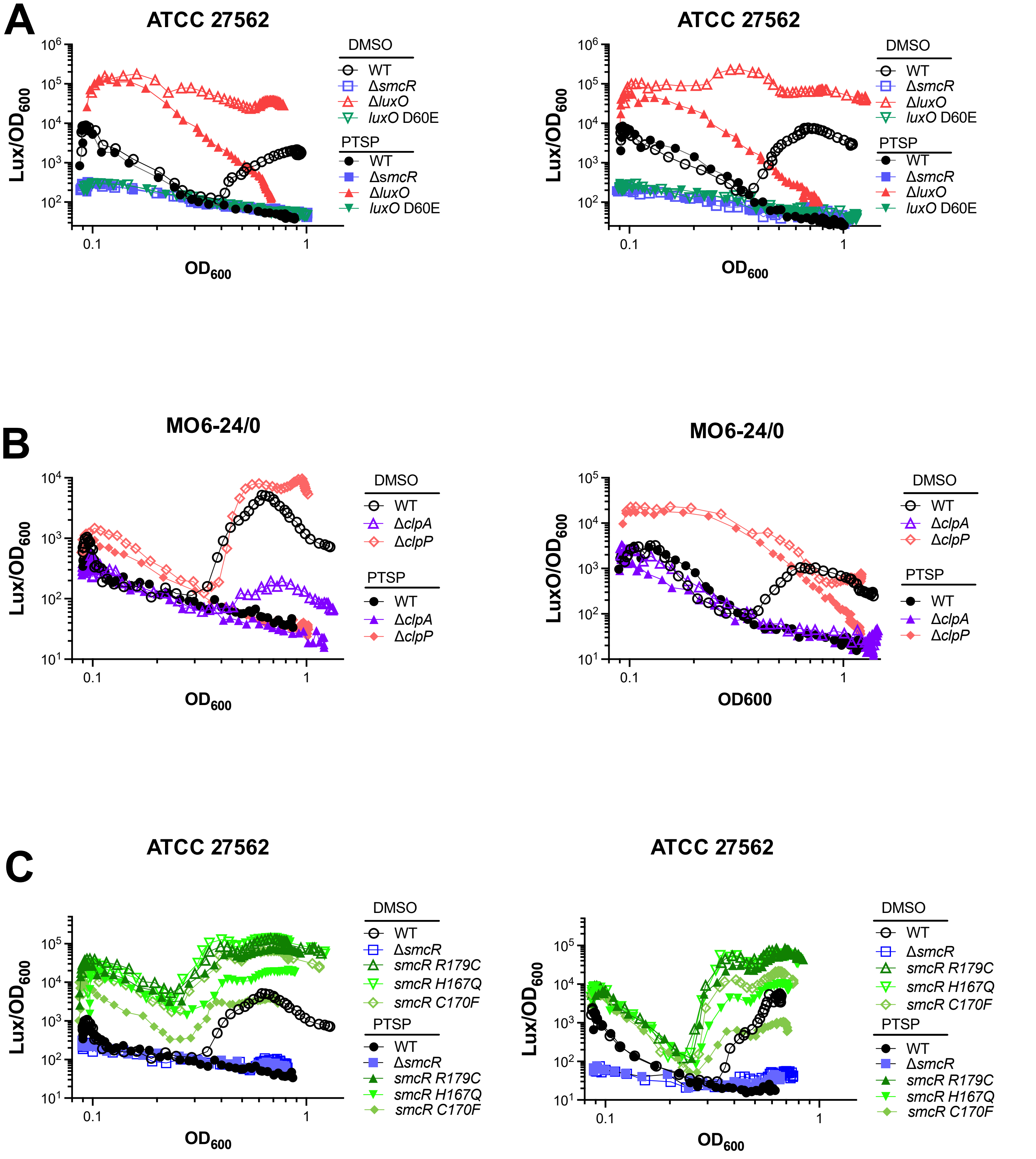

**Figure S15. ClpAP proteolysis of SmcR is blocked by substitutions that block inhibitor binding.** (A, B, C) Bioluminescence assays are plotted (Lux/$OD_{600}$) for *V. vulnificus* strains ATCC 27562 (A, C) and MO6-24/0 (B) and isogenic mutants. Strains were treated with 25 μM PTSP (+) or an equivalent volume of DMSO solvent (-). Each panel shows two biological replicates; the third biological replicate is in the main figure.

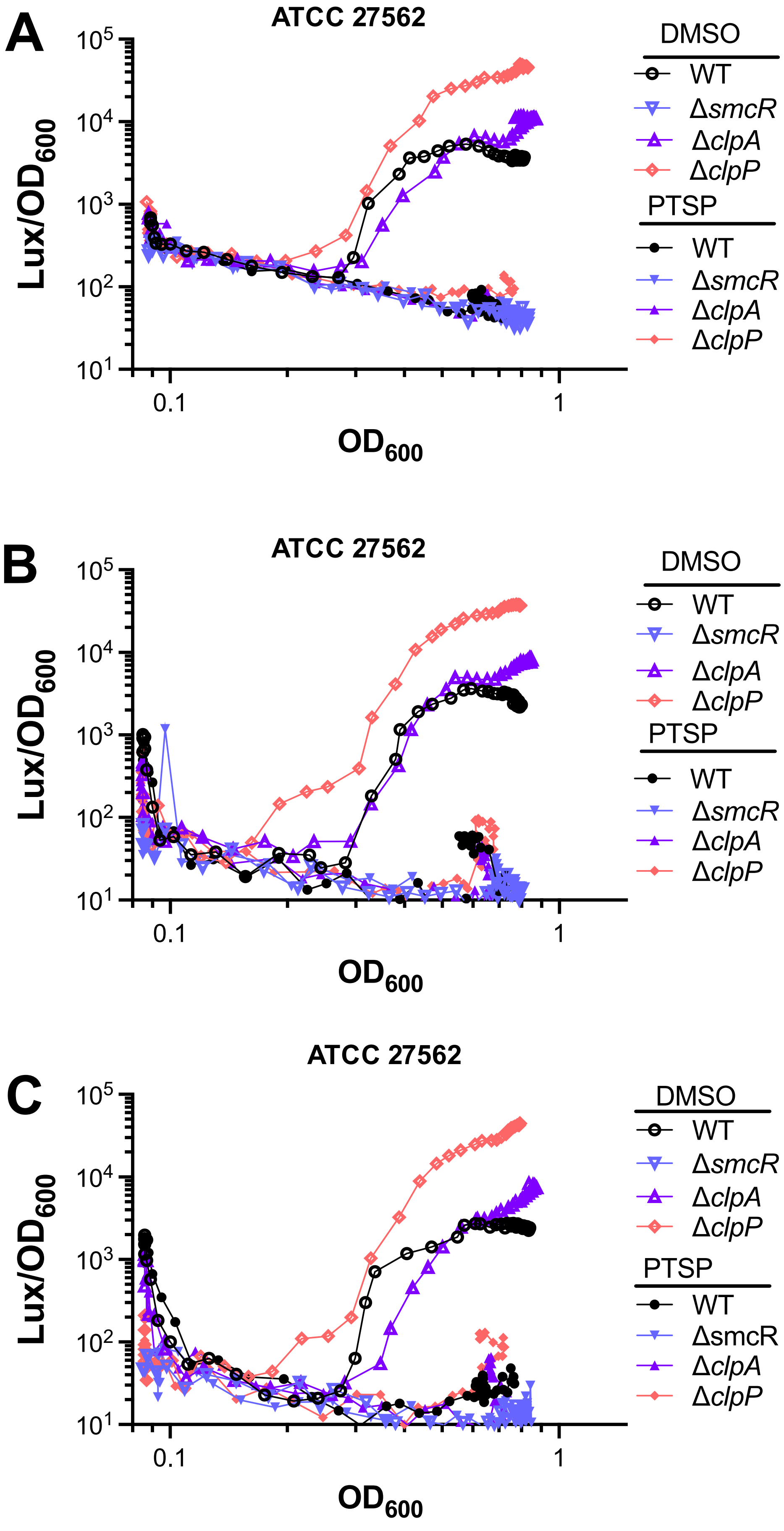

**Figure S16. ClpAP proteolysis of SmcR is blocked by substitutions that block inhibitor binding.** Bioluminescence assays are plotted (Lux/$OD_{600}$) for *V. vulnificus* strain ATCC 27562 and isogenic mutants. Strains were treated with 25 μM PTSP (+) or an equivalent volume of DMSO solvent (-). Each panel shows a biological replicate.
